# Supervised-learning is an accurate method for network-based gene classification

**DOI:** 10.1101/721423

**Authors:** Renming Liu, Christopher A Mancuso, Anna Yannakopoulos, Kayla A Johnson, Arjun Krishnan

## Abstract

**Background:** Assigning every human gene to specific functions, diseases, and traits is a grand challenge in modern genetics. Key to addressing this challenge are computational methods such as supervised-learning and label-propagation that can leverage molecular interaction networks to predict gene attributes. In spite of being a popular machine learning technique across fields, supervised-learning has been applied only in a few network-based studies for predicting pathway-, phenotype-, or disease-associated genes. It is unknown how supervised-learning broadly performs across different networks and diverse gene classification tasks, and how it compares to label-propagation, the widely-benchmarked canonical approach for this problem.

**Results:** In this study, we present a comprehensive benchmarking of supervised-learning for network-based gene classification, evaluating this approach and a state-of-the-art label-propagation technique on hundreds of diverse prediction tasks and multiple networks using stringent evaluation schemes. We demonstrate that supervised-learning on a gene’s full network connectivity outperforms label-propagation and achieves high prediction accuracy by efficiently capturing local network properties, rivaling label-propagation’s appeal for naturally using network topology. We further show that supervised-learning on the full network is also superior to learning on node-embeddings (derived using *node2vec*), an increasingly popular approach for concisely representing network connectivity.

**Conclusion:** These results show that supervised-learning is an accurate approach for prioritizing genes associated with diverse functions, diseases, and traits and should be considered a staple of network-based gene classification workflows. The datasets and the code used to reproduce the results and add new gene classification methods have been made freely available.

## Introduction

In the post-genomic era, a grand challenge is to characterize every gene across the genome in terms of the cellular pathways they participate in, and which multifactorial traits and diseases they are associated with. Computationally predicting the association of genes to pathways, traits, or diseases – the task termed here as “gene classification” – has been critical to this quest, helping prioritize candidates for experimental verification and for shedding light on poorly characterized genes [1–7]. Key to the success of these methods has been the steady accumulation of large amounts of publicly available data collections such as curated databases of genes and their various attributes [8–18], controlled vocabularies of biological terms organized into ontologies [19–22], high-throughput functional genomic assays [23–25], and molecular interaction networks [26–30].

While protein sequence and 3D structure are remarkably informative about the corresponding gene’s molecular function [3,7,31–33], the pathways or phenotypes that a gene might participate in significantly depends on the other genes that it works with in a context dependent manner.. Molecular interaction networks – graphs with genes or proteins as nodes and their physical or functional relationships as edges – are powerful models for capturing the functional neighborhood of genes on a whole-genome scale [34–36]. These networks are often constructed by aggregating multiple sources of information about gene interactions in a context-specific manner [29,37]. Therefore, unsurprisingly, several studies have taken advantage of these graphs to perform network-based gene classification [38–43].

The canonical principle of network-based gene classification is “guilt-by-association”, the notion that proteins/genes that are strongly connected to each other in the network are likely to perform the same functions, and hence, participate in similar higher-level attributes such as phenotypes and diseases [44]. Instead of just aggregating “local” information from direct neighbors [45], this principle is better realized by propagating pathway or disease labels across the network to capture “global” patterns, achieving state-of-the-art results [34–36,38–41,46–56]. These global approaches belong to a class of methods referred to here as “label propagation.” Distinct from label propagation is another class of methods for gene classification that relies on the idea that network patterns characteristic of genes associated with a specific phenotype or pathway can be captured using supervised machine learning [29,42,43,57–59]. While this class of methods – referred to here as “supervised learning” – has yielded promising results in a number of applications, how it broadly performs across different types of networks and diverse gene classification tasks is unknown. Consequently, supervised learning is used far less compared to label propagation for network-based gene classification.

The goal of this study is to perform a comprehensive, systematic benchmarking of supervised learning (SL) for network-based gene classification across a number of genome-wide molecular networks and hundreds of diverse prediction tasks using meaningful evaluation schemes. Within this rigorous framework, we compare supervised learning to a widely-used, state-of-the-art label propagation (LP) technique, testing both the original (adjacency matrix A) and a diffusion-based representation of the network (influence matrix I; Fig. 1). This combination results in four methods (listed with their earliest known references): label-propagation on the adjacency matrix (LP-A) [45], label-propagation on the influence matrix (LP-I) [52], supervised-learning on the adjacency matrix (SL-A) [58], and supervised-learning on the influence matrix (SL-I) [57]. Additionally, we evaluate the performance of supervised learning using node embeddings as features, as the use of node embeddings is burgeoning in network biology.

**Fig. 1.**
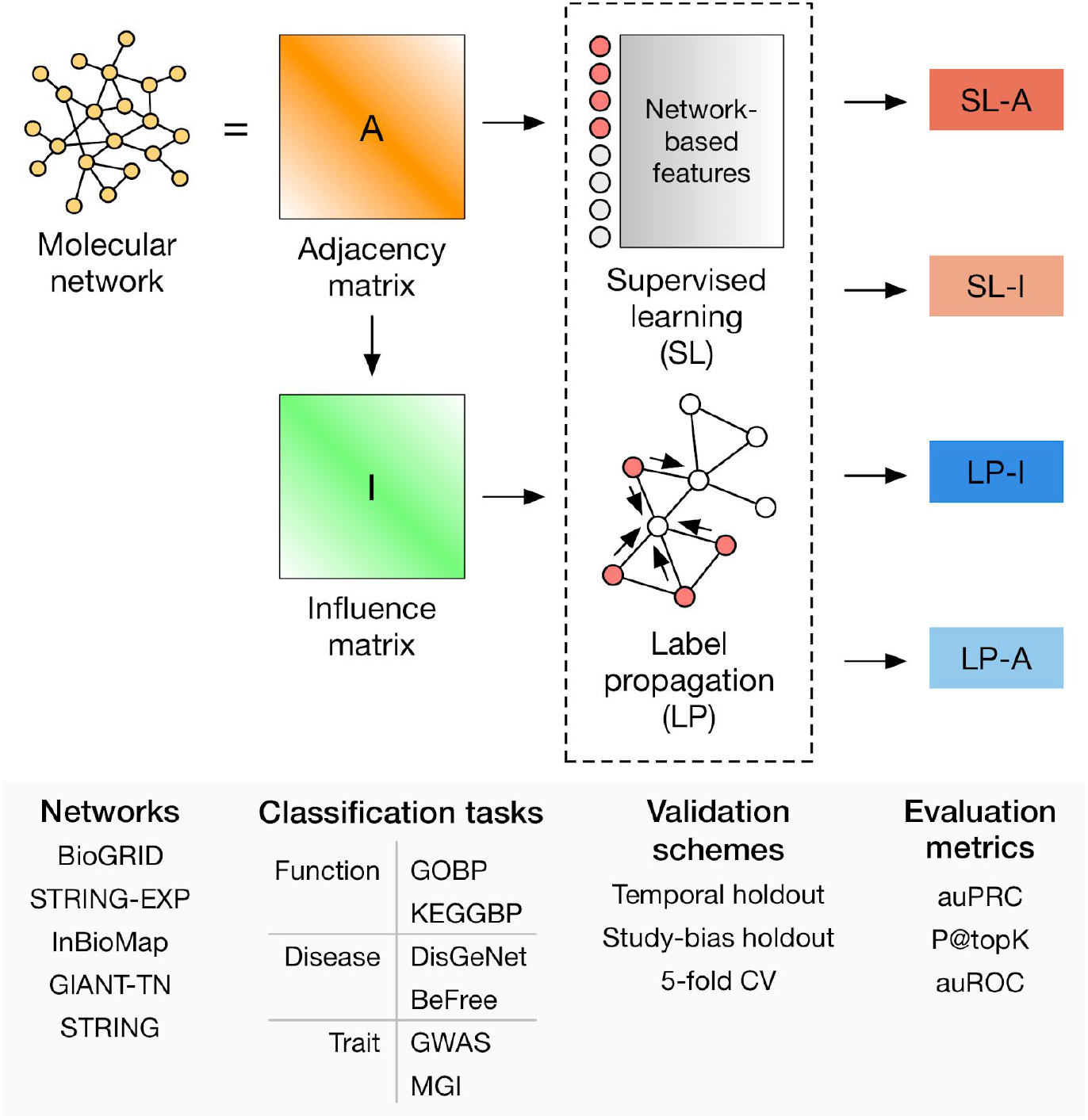
Workflow for gene classification pipeline. Four methods are compared: supervised learning on the adjacency matrix (SL-A), supervised learning on the influence matrix (SL-I), label propagation on the adjacency matrix (LP-A), and label propagation on the influence matrix (LP-I). Model performance on a variety of gene classification tasks is evaluated over a number of different molecular networks, validation schemes, and evaluation metrics. Additionally, the performance of supervised learning using node embeddings as features (SL-E) is evaluated (not shown in this figure).

Our results demonstrate that SL outperforms LP for gene-function, gene-disease, and gene-trait prediction. We also observe that SL captures local network properties as efficiently as LP, where both methods achieve more accurate predictions for genesets that are more tightly clustered in the network. Lastly, we show that SL using the full network connectivity is superior to using low-dimensional node embeddings as the features, which, in turn, is competitive to LP.

## Methods and Data

### Networks

We chose a diverse set of undirected, human gene/protein networks based on criteria laid out in [30] (Fig. 1): 1) networks constructed using high- or low-throughput data, 2) the type of interactions the network was constructed from, and 3) if annotations were directly incorporated in constructing the network. We used versions of the networks that were released prior to 2017 so as not to bias the temporal holdout evaluations. We used all edge scores (weights) unless otherwise noted, and the nodes in all networks were mapped into Entrez genes using the MyGene.info database [16,17]. If the original node ID mapped to multiple Entrez IDs, we added edges between all possible mappings. The networks used in this study are BioGRID [28], the full STRING network [26] as well as the subset with just experimental support (referred to as STRING-EXP in this study), InBioMap [27], and the tissue-naïve network from GIANT [29], referred to as GIANT-TN in this study. These networks cover a wide size range, with the number of nodes ranging from 14,089 to 25,689 and the number of edges ranging from 141,629 to 38,904,929. More information on the networks can be found in the Supplemental Material (Section 1.1).

### Network Representations

We considered three distinct representations of molecular networks: the adjacency matrix, an influence matrix, and low-dimensional node embeddings. Let *G* = (*V*, *E*, *W*) denote an undirected molecular network, where *V* is the set of vertices (genes), *E* is the set of edges (associations between genes), and *W* is the set of edge weights (the strengths of the associations). *G* can be represented as a weighted adjacency matrix *A_i,j_* = *W_i,j_*, where *A* ∈ *R*^|*V*|×|*V*|^. *G* can also be represented as an influence matrix, *F* ∈ *R*^|*V*|×|*V*|^, which can capture both local and global structure of the network. *F* was obtained using a random walk with restart transformation kernel [41],

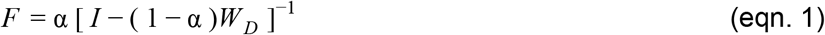

where, α is the restart parameter, *I* is the identity matrix, and *W_D_* is the degree weighted adjacency matrix given by *W_D_* = *AD*^−1^, where *D* ∈ *R*^|*V*|*x*|*V*|^ is a diagonal matrix of node degrees. A restart parameter of 0.85 was used for every network in this study.

*G* can also be transformed into a low-dimensional representation through the process of node embedding. In this study we used the *node2vec* algorithm [60], which borrows ideas from the *word2vec* algorithm [61,62] from natural language processing. The objective of *node2vec* is to find a low-dimensional representation of the adjacency matrix, *E* ∈ *R*^|*V*|×*d*^, where *d* ≪ |*V*|. This is done by optimizing the following log-probability objective function:

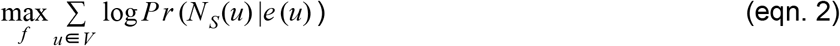

where *N_S_*(*u*)is the network neighborhood of node *u* generated through a sampling strategy *S*, and *e*(*u*) ∈ *R^d^* is the feature vector of node *u*. In *node2vec*, the sampling strategy is based on random walks that are controlled using two parameters *p* and *q*, in which a high value of *q* keeps the walk local (a breadth-first search), and a high value of *p* encourages outward exploration (a depth-first search). The values of *p* and *q* were both set to 0.1 for every network in this study.

### Prediction methods

We compared the prediction performance across four specific methods across two classes, label-propagation (LP) and supervised-learning (SL).

#### Label Propagation

LP methods are the most widely used methods in network-based gene classification and achieve state-of-the-art results [39,48]. In this study, we considered two LP methods, label propagation on the adjacency matrix (LP-A) and label propagation on the influence matrix (LP-I). First, we constructed a binary vector of ground-truth labels, *x* ∈ *R*^|*V*|×1^, where *x_i_* = 1 if gene *i* is a positively labeled gene in the training set, and 0 otherwise. In LP-A, we constructed a score vector, *S* ∈ *R*^|*V*|×1^, denoting the predictions,

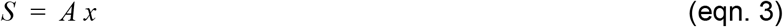

where *A* is the adjacency matrix. Thus, the predicted score for a gene using LP-A is equal to the sum of the weights of the edges between the gene and its direct, positively labeled network neighbors. In LP-I, the score vector, *S*, is generated using eqn. 3, except *A* is by replaced by *F*, the influence matrix (eqn. 1). In both LP-A and LP-I, only positive examples in the training set are used to calculate the score vector *S*, but both positive and negative examples in the test set are later used for evaluation.

#### Supervised Learning

Supervised-learning (SL) can be used for network-based gene classification by using each gene’s network neighborhoods as feature vectors, along with gene labels, in a classification algorithm. Here, we used logistic regression with L2 regularization as the SL classification algorithm, which is a linear model that aims to minimize the following cost function [63]:

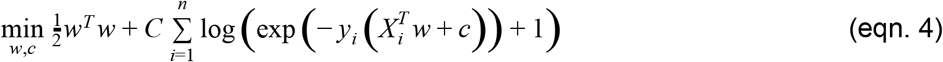

where *w* ∈ *R^m^* is the vector of weights for a model with *m* features, *C* determines the regularization strength, *n* the number of samples, *y* is the ground-truth label, *X* ∈ *R*^*n*×*m*^ is the data matrix, and *c* is the intercept. After training a model using the labeled genes in the training set, the learned model weights are used to classify the genes in the testing set, returning a prediction probability for these genes that is bounded between 0 and 1. The regularization parameter, *C*, was set to 1.0 for all models in this study.

In this study, three different network-based gene-level feature vectors were used to train three different SL classifiers: the rows of the adjacency matrix (SL-A), the rows of the influence matrix (SL-I), and the rows of the node embedding matrix (SL-E). Model selection and hyperparameter tuning are described in detail in the Supplemental Material (Section 1.2).

### Geneset-collections

We curated a number of geneset-collections to test predictions on a diverse set of tasks: function, disease, and trait (Fig. 1). Function prediction was defined as predicting genes associated with biological processes that are part of the Gene Ontology (referred to here as “GOBP”) [19,20] obtained from MyGene.info [16,17] and pathways from the Kyoto Encyclopedia of Genes and Genomes [10–12], referred to “KEGGBP” since disease-related pathways were removed from the original KEGG annotations in the Molecular Signatures Database [8,9]. Disease prediction was defined based on predicting genes associated with diseases in the DisGeNET database [13,14]. Annotations from this database were divided into two separate geneset-collections: those that were manually-curated (referred to as “DisGeNet” in this study) and those derived using the BeFree text-mining tool (referred to as “BeFree” in this study). Trait prediction was defined as predicting genes linked to human traits from Genome-wide Association Studies (GWAS), curated from a community challenge [64], and mammalian phenotypes (annotated to human genes) from the Mouse Gene Informatics (MGI) database [15].

Each of these six geneset-collections contained anywhere from about a hundred to tens of thousands of genesets that varied widely in specificity and redundancy. Therefore, each collection was preprocessed to ensure that the final set of prediction tasks from each source are specific, largely non-overlapping, and not driven by multi-attribute genes. First, if genesets in a collection corresponded to terms in an ontology (e.g. biological processes in the GOBP collection), annotations were propagated along the ontology structure to obtain a complete set of annotations for all genesets. Second, we removed genesets if the number of genes annotated to the geneset was above a certain threshold and then compared these genesets to each other in order to remove genesets that were highly-overlapping with other genesets in the collection, resulting in a set of specific, non-redundant genesets. Finally, individual genes that appeared in more than 10 of the remaining genesets in a collection were removed from all the genesets in that collection to remove multi-attribute (e.g. multi-functional) genes that are potentially easy to predict [65]. Detailed information on geneset pre-processing and geneset attributes can be found in the Supplemental Material (Section 1.3, Table S2, and Fig. S3).

#### Selecting positive and negative examples

In each geneset collection, for a given geneset, genes annotated to that set were designated as the set of positive examples. The SL methods additionally required a set of negative genes for each given geneset for training; both SL and LP methods require a set of negative genes for each geneset for testing. A set of negative genes was generated by: a) finding the union of all genes annotated to all genesets in the collection, b) removing genes annotated to the given geneset, and c) removing genes annotated to any geneset in the collection which significantly overlapped with the given geneset (p-value < 0.05 based on the one-sided Fisher’s exact test).

### Validation schemes

We performed extensive and rigorous evaluations based on three validation schemes: temporal holdout, study-bias holdout, and 5-fold cross validation (5FCV). In temporal holdout, within a geneset-collection, genes that only had an annotation to any geneset in the collection after Jan 1st, 2017 were considered test genes, and all other genes were considered training genes. Temporal holdout is the most stringent evaluation scheme for gene classification since it mimics the practical scenario of using current knowledge to predict the future and is the preferred evaluation method used in the CAFA challenges [3,7]. Since Gene Ontology was the only source with clear date-stamps for all its annotations, temporal holdout was applied only to the GOBP geneset-collection. For study-bias holdout, genes were ranked by the number of PubMed articles they were mentioned in, obtained from [66]. The top two-thirds of the most-mentioned genes were considered training genes, and the rest of the least-mentioned genes were used for testing. Study-bias holdout mimics the real-world situation of learning from well-characterized genes to predict novel un(der)-characterized genes. The last validation scheme is the traditional 5-fold cross validation, where the genes are split into 5 equal folds in a stratified manner (i.e. in each split, the proportion of genes in the positive and negative classes is preserved). In all these schemes, only genesets with at least 10 positive genes in both the training and test sets were considered. More information on the validation schemes is available in the Supplemental Material (Section 1.4).

### Evaluation Metrics

In this study, we considered three evaluation metrics: the area under the precision-recall curve (auPRC), the precision of the top *K* ranked predictions (P@TopK), and, for completeness, the area under the receiver-operator curve (auROC). For P@TopK, we set *K* equal to the number of ground truth positives in the testing set. Since the standard auPRC and P@TopK scores are influenced by the prior probability of finding a positive example (equal to the proportion of positives to the total of positives and negatives), we expressed both metrics as the logarithm (base 2) of the ratio of the original metric to the prior. More details on the evaluation metrics can be found in the Supplemental Material (Section 1.5).

## Results

We systematically compare the performance of four gene classification methods (Fig. 1): supervised learning on the adjacency matrix (SL-A), supervised learning on the influence matrix (SL-I), label propagation on the adjacency matrix (LP-A), and label propagation on the influence matrix (LP-I). We choose six geneset-collections that represent three prominent gene-classification tasks: gene-function (GOBP, KEGGBP), gene-disease (DisGeNet, BeFree), and gene-trait (GWAS, MGI) prediction. We use three different validation schemes: temporal holdout (train on genes annotated before 2017 and test on genes annotated in 2017 or later; only done for GOBP as it has clear timestamps), holdout based on study-bias (train on well-studied genes and predict on less-studied genes), and the traditional 5-fold cross validation (5FCV). Temporal holdout and study-bias holdout validation schemes are presented in the main text as they are more stringent and reflective of real-world tasks as compared to 5FCV [67]. To ascertain the robustness of the relative performance of the methods to the underlying network, we choose five different genome-scale molecular networks that differ in their content and construction. To be in concert with temporal holdout evaluation and curtail data leakage, all the networks used throughout this study are the latest versions released before 2017. We present evaluation results based on the area under the precision-recall curve (auPRC) in the main text and results based on the precision at top-k (P@topK) and the area under the ROC curve (auROC) in the Supplemental Material (Figs. S4-S8). We note that the 5FCV, P@topK, and auROC results in the Supplemental Material are, for the most part, consistent with the results presented in the main text of this study.

Our first analysis was to directly compare all four prediction methods against each other for each geneset in a given collection. For each geneset-collection–network combination, we rank the four methods per geneset (based on auPRC) using the standard competition ranking and calculate each method’s average rank across all the genesets in the collection (Fig. 2). For function prediction, SL-A is the top-performing method by a wide margin (particularly clear based on GOBP temporal holdout), with SL-I being the second best method. For disease and trait prediction, SL-A and SL-I still outperform LP-I, but to a lesser extent. In all cases, LP-A is the worst performing method. The large performance difference between the SL and LP methods in the GOBP temporal holdout validation is noteworthy since temporal holdout is the most stringent validation scheme and the one employed in community challenges such as CAFA [3,7].

**Fig. 2.**
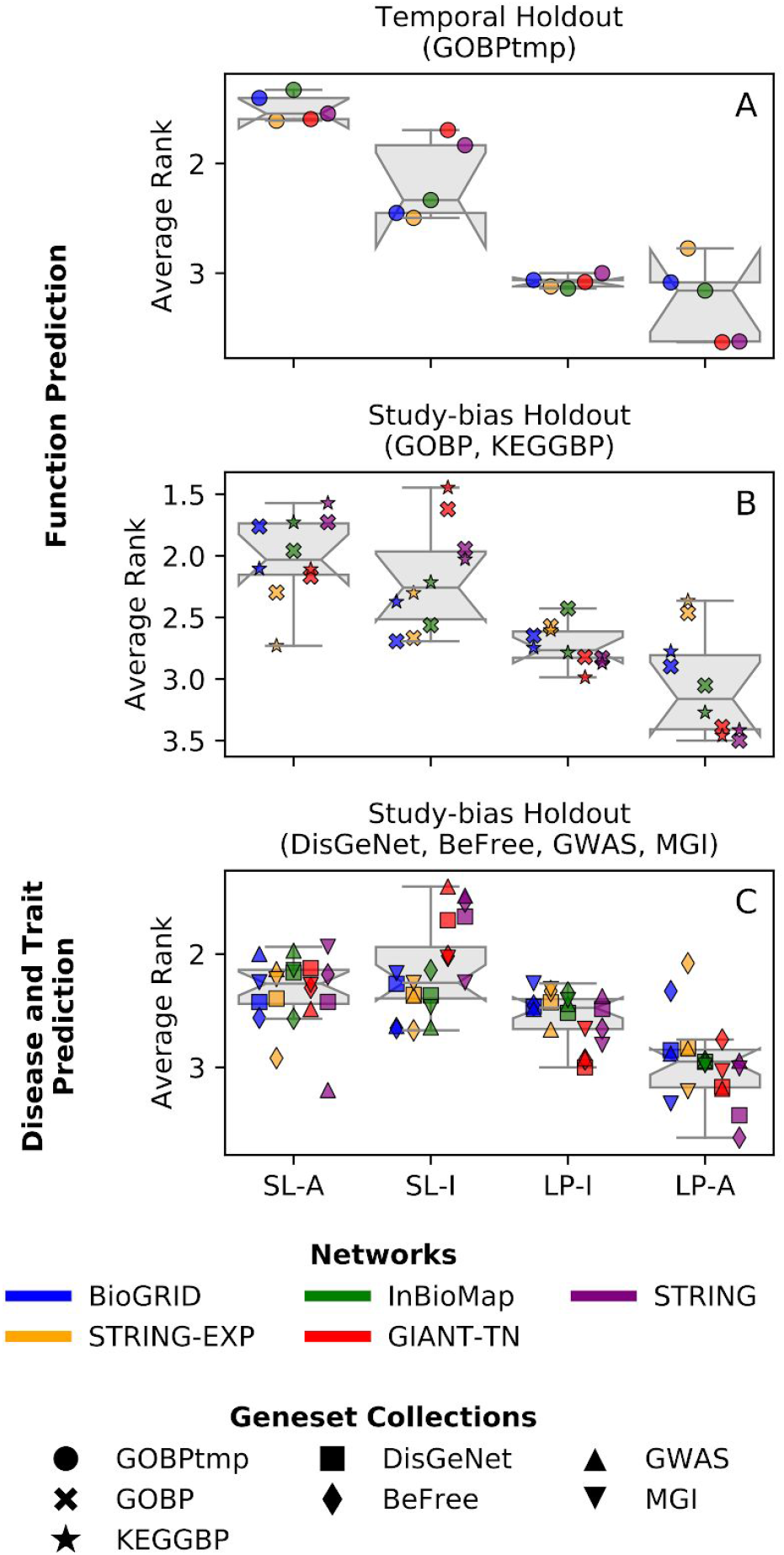
Average rank across the four methods. Each point in each boxplot represents the average rank for a geneset-collection–network combination, obtained based on ranking the four methods in terms of performance for each geneset in a geneset-collection using the standard competition ranking. (A) Functional prediction tasks using GOBP temporal holdout, (B) Functional prediction tasks using study-bias holdout for GOBP and KEGGBP, and (C) Disease and trait prediction tasks using study-bias holdout for DisGeNet, BeFree, GWAS, and MGI. The results are shown for auPRC where different colors represent different networks and different marker styles represent the different geneset-collections. SL methods outperform LP methods for all prediction tasks.

Following the observation that SL methods outperform LP methods based on relative ranking, we use a non-parametric paired test (Wicoxon signed-rank test) to statistically assess the difference between specific pairs of methods (Fig. 3A). For each geneset-collection–network combination, we compare the two methods in one class to the two methods in the other class (i.e. we compare SL-A to LP-A, SL-A to LP-I, SL-I to LP-A, and SL-I to LP-I). Each comparison yields a p-value along with the number of genesets in the collection where one method outperforms the other. After correcting the four p-values for multiple hypothesis testing [68], if a method from one class outperforms both methods from the other class independently (in terms of the number winning genesets), and if both (corrected) p-values are <0.05, we consider a method to have significantly better performance compared to the entire other class. Additionally, we track the percentage of times the SL methods outperform the LP methods across all four comparisons within a geneset-collection–network combination.

**Fig. 3.**
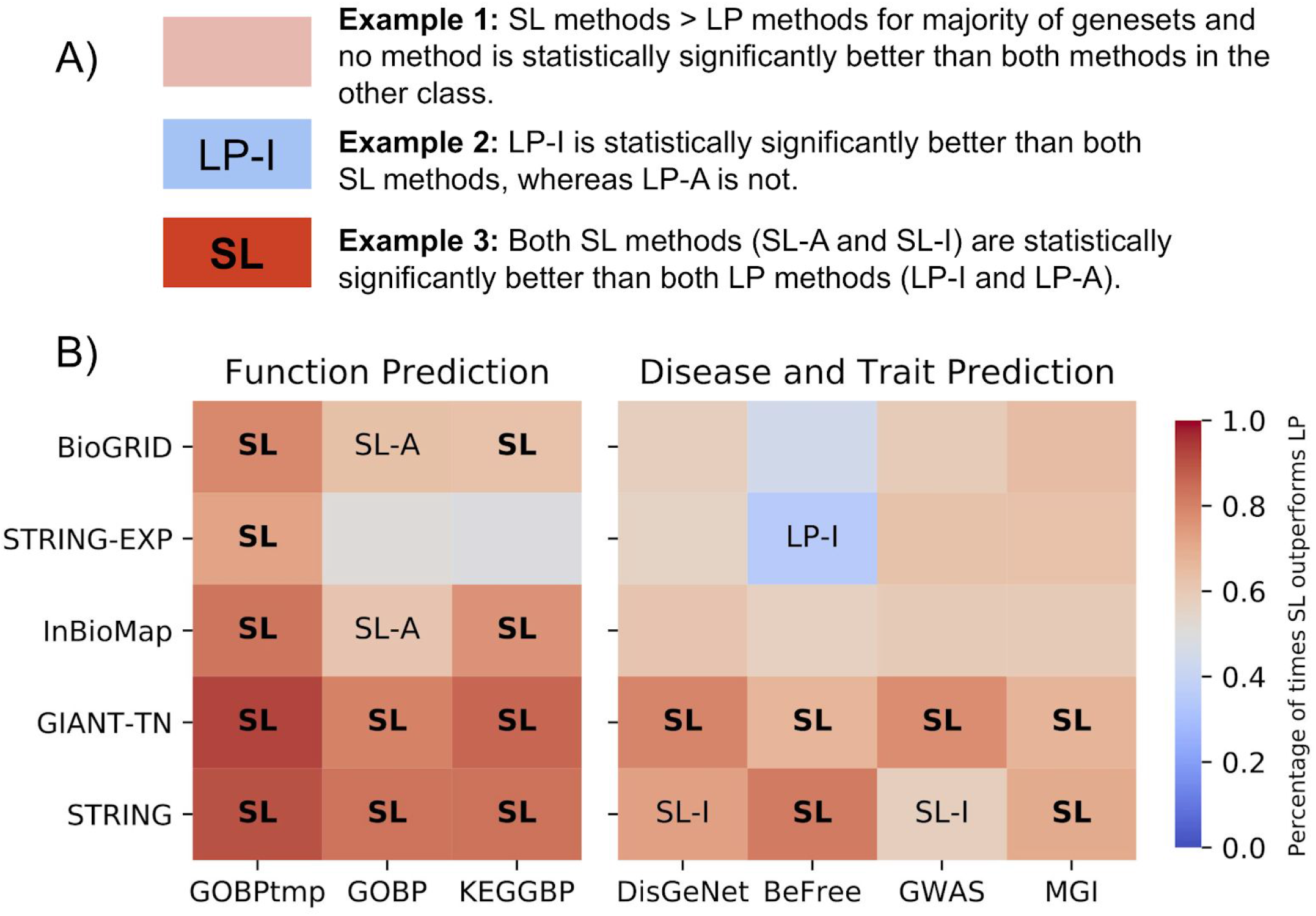
Testing for a statistically significant difference between SL and LP methods. A) A key on interpreting the analysis. For each network-geneset combination, each method is compared to the two methods from the other class (i.e. SL-A vs LP-I, SL-A vs LP-A, SL-I vs LP-I, SL-I vs LP-A). If a method was found to be significantly better than both methods from the other class (Wilcoxon ranked-sum test with an FDR threshold of 0.05), the cell is annotated with that method. If both models in that class were found to be significantly better than the two methods in the other class, the cell is annotated in bold with just the class. The color scale represents the fraction of genesets that were higher for the SL methods across all four comparisons. The first column uses GOBP temporal holdout, whereas the remaining 6 columns use study-bias holdout. B) SL methods show a statistically significant improvement over LP methods, especially for function prediction.

The results show that, for function prediction, SL is almost always significantly better than LP when considering auPRC (Fig. 3B). Based on temporal holdout on GOBP, both SL-A and SL-I are always significantly better than both LP methods. Based on study-bias holdout, in the 10 function prediction geneset-collections–network combinations using GOBP and KEGGBP, SL-A is a significantly better method 8 times (80%) and SL-I is a significantly better method 6 times (60%). Neither LP-I nor LP-A ever significantly outperform the SL models. The performance of SL and LP are more comparable for disease and trait prediction, but SL methods still perform better in a larger fraction of genesets. For the 20 disease and trait geneset-collection–network combinations, SL-I is a significantly better method 8 times (40%), and SL-A is a significantly better method 6 times (30%), LP-I is a significantly better method once (5%), and LP-A is never a significantly better method.

To visually inspect not only the relative performance of all four methods, but to also see how well the models are performing in an absolute sense, we examined the boxplots of the auPRC values for every geneset-collection–network combination (Fig. 4). The first notable observation is that, regardless of the method, function prediction tasks have much better performance results than disease/trait prediction tasks (Fig. 4B). Based on temporal holdout for function prediction (GOBPtmp), SL-A is the top-performing model based on the highest median performance for every network. Additionally, for all networks except STRING-EXP, SL-I is the second best performing model. For the 10 combinations of five networks with GOBP and KEGGBP, the top method based on the highest median performance is an SL method all but once, with SL-A being the top model 7 times (70%), SL-I being the top model 2 times (20%, GOBP and KEGGBP on GIANT-TN), and LP-A being the top model once (10%, KEGGBP on STRING-EXP). As noted earlier, for disease and trait prediction, SL and LP methods have more comparable performance. Of the 20 geneset-collection–network combinations, each of SL-A, SL-I, LP-I, and LP-A is the top method based on median performance 5 (25%), 10 (50%), 4 (20%), and 1 (5%) times, respectively.

**Fig. 4.**
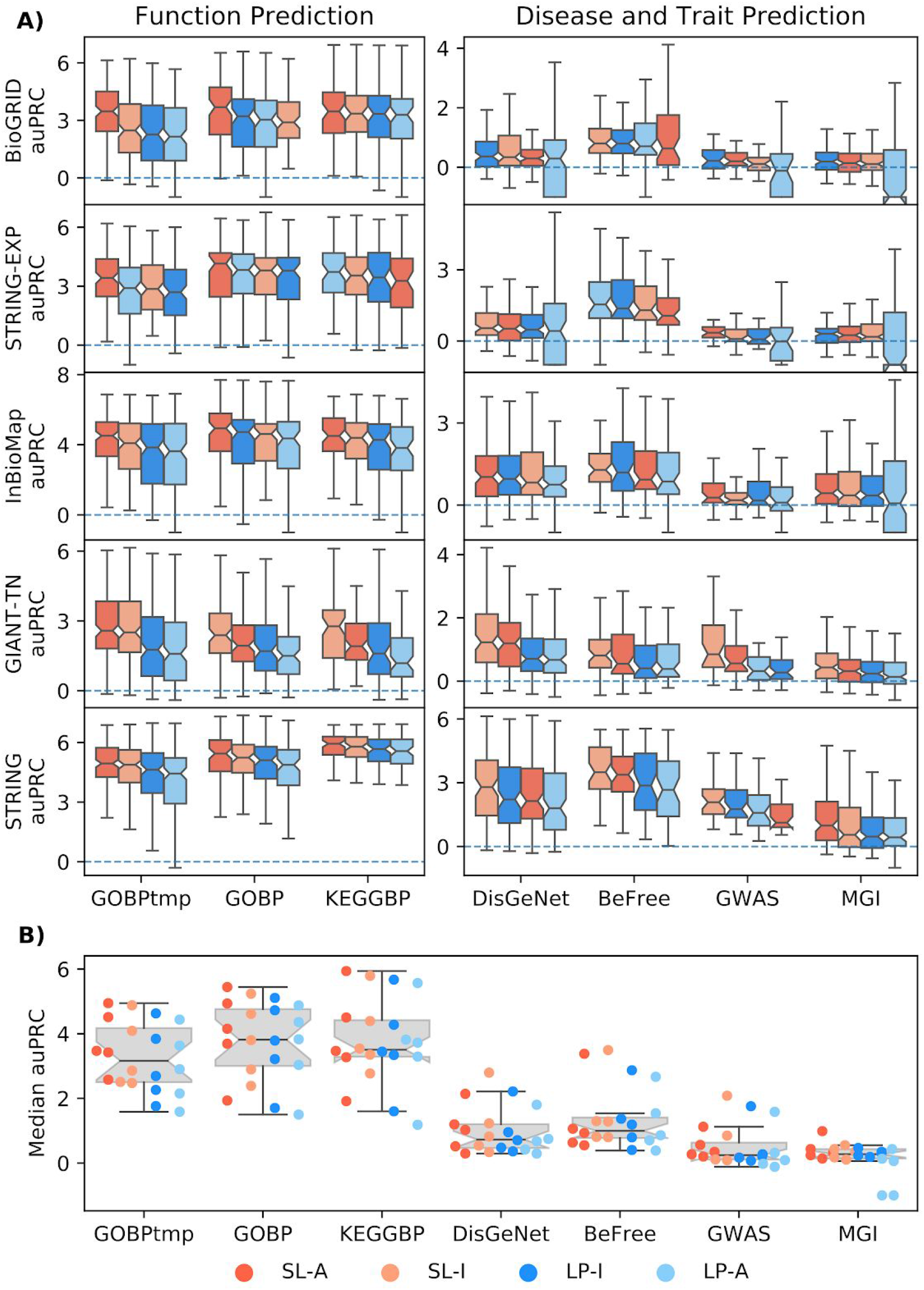
Boxplots for performance across all geneset-collection–network combinations. A)The performance for each individual geneset-collection–network combination is compared across the four methods; SL-A (red), SL-I (light red), LP-I (blue), and LP-A (light blue). The methods are ranked by median value with the highest scoring method on the left. Results show SL methods outperform LP methods, especially for function prediction. B) Each point in the plot is the median value from one of the boxplots in A. This shows that both SL and LP methods perform better for function prediction compared to disease/trait prediction.

Among the two classes of network-based models – SL and LP – it is intuitively clear how LP directly uses network connections to propagate information from the positively-labeled nodes to other nodes close in the network. On the other hand, while SL is an accurate method for gene classification, it has not been studied if SL’s performance is tied to any traditional notion of network connectivity. To shed light on this problem, we investigated the performance of SL-A and LP-I as a function of three different properties of individual genesets in a collection: the number of annotated genes, edge density (a measure of how tightly connected the geneset is within itself), and segregation (a measure of how isolated the geneset is from the rest of the network). While the performance of neither SL-A nor LP-I has a strong association with the size of the geneset, the performance of SL-A has a strong positive correlation with both edge density and segregation of the geneset, similar to what is seen for LP-I (Fig. 5). For visual clarity, Fig. 5 presents results for just the STRING network, but very similar results are seen in the other networks as well (Fig. S9). Detailed information on how the geneset and network properties are calculated can be found in the Supplemental Material (Section 1.3).

**Fig. 5.**
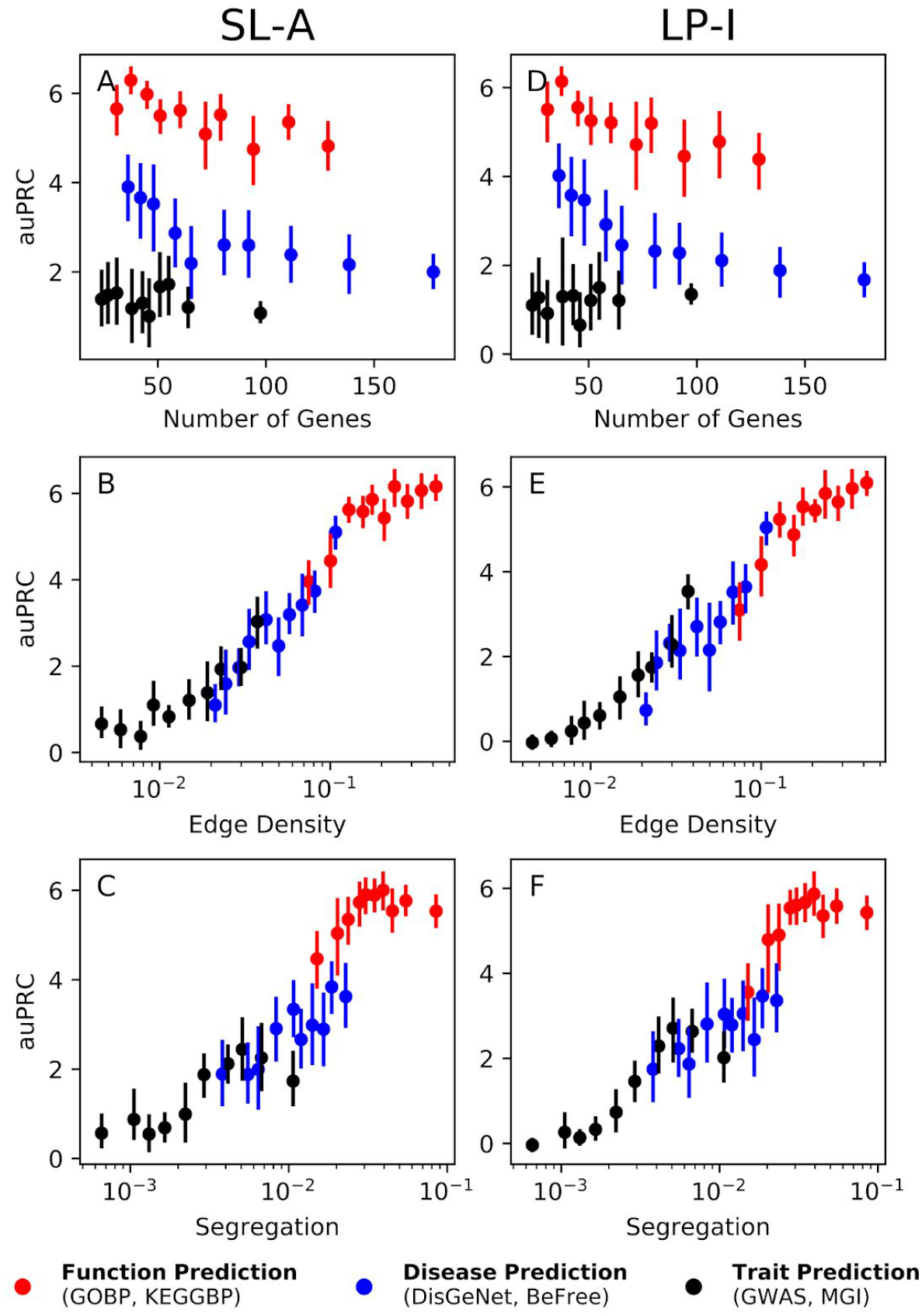
Performance vs Network/Geneset properties. SL-A (A-C) is able to capture network information as efficiently as LP-I (D-F), for the STRING network. There is no correlation between the number of genes in the geneset versus performance (A,D), but there is a strong correlation between the performance and the edge density (B,E) as well as segregation (C,F). The different colored dots represent function genesets (red, GOBP and KEGGBP), disease genesets (blue, DIGenet and BeFree), and trait genesets (black, GWAS and MGI). The vertical line is the 95% confidence interval. Similar trends can be seen for the other networks (Fig. S9).

Finally, since machine learning on node embeddings is gaining popularity for network-based node classification, we compare the top SL and LP methods tested here to this approach. Specifically, we compare LP-I and SL-A to an SL method using embeddings (SL-E) obtained from the *node2vec* algorithm [60] (Fig. 6). For function prediction, we observe that SL-E substantially outperforms LP-I. For GOBP temporal holdout, SL-E is always significantly better than LP-I. For the GOBP and KEGGBP study-bias holdout, out of the 10 geneset-collection–network combinations, SL-E is significantly better than LP-I 5 times (50%), whereas the converse is true only once (10%). These patterns nearly reverse for the 20 disease/trait prediction tasks, with LP-I performing significantly better than SL-E 6 times (30%), and SL-E significantly outperforming LP-I 3 times (15%). The comparison between SL-E and SL-A showed that SL-A demonstrably outperforms SL-E for both function and disease/trait prediction tasks. Among the 30 geneset-collection–network combinations, SL-A is a significantly better model 20 times (67%), whereas SL-E comes out on top just once (3%). This shows that although methods that use node embeddings are a promising avenue of research, they should be compared to the strong baseline of SL-A when possible.

**Fig. 6.**
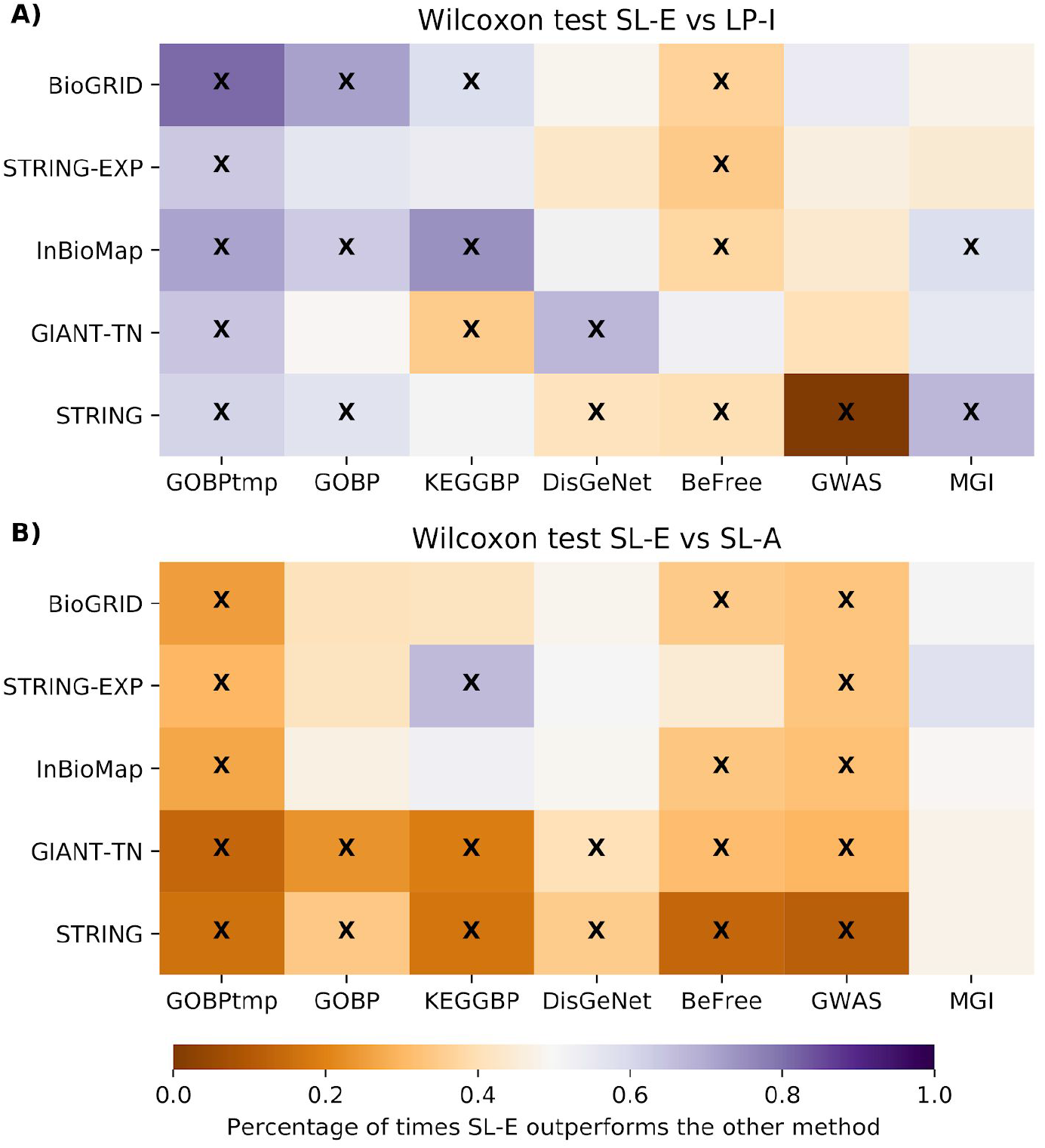
Performance of SL-E vs LP-I and SL-A. We compare the performance of supervised learning on the embedding matrix (SL-E) vs LP-I and SL-A using a Wilcoxon ranked-sum test. The performance metric is auPRC, the color scale represents the fraction of terms that were higher for the SL-E model (with purple being SL-E had a higher fraction of better performing genesets compared to either LP-I or SL-A) and an “x” signifies that the p-value from the Wilcoxon test was below 0.05. A) Shows that SL-E is quite competitive with the current state-of-the-art method of LP-I and B) shows that SL-A outperforms SL-E in a majority of cases.

## Discussion

We have conducted the first comprehensive benchmarking of supervised-learning that establishes it as a leading approach for network-based gene classification. Further, to the best of our knowledge, neither the studies that propose new methods nor those that systematically compare existing approaches have directly compared the two classes of methods – supervised-learning and label-propagation – against each other. Our work, provides this systematic comparison and shows that supervised-learning (SL) methods demonstrably outperform label-propagation (LP) methods for network-based gene classification, particularly for function prediction.

Both SL and LP methods are, in general, more accurate for function prediction than disease and trait prediction. This trend is likely due to the fact that molecular interaction networks are primarily intended, either through curation or reconstruction, to reflect biological relationships between genes/proteins as they pertain to ‘normal’ cellular function. The utility of network connectivity to gene-disease or gene-trait prediction is incidental to the information the network holds about gene-function associations. This notion is supported by the observation that genesets related to function genesets are more tightly-clustered than disease and trait genesets in the genome-wide molecular networks used in this study (Fig. S3). Further analysis of prediction accuracy of genesets as a function of their network connectivity lends credence to the use of network structure by SL (Fig. 5 and Fig. S9). Part of LP’s appeal, widespread use, and development is this natural use of network topology to predict gene properties by diffusing information from characterized genes to uncharacterized genes in their network vicinity. Therefore, we expect that genes associated with tightly-clustered pathways, traits, or diseases will be easier to predict using LP, which is observed in our analysis (Fig. 5 and Fig. S9). On the other hand, since SL (based on the full network) is designed to use global gene connectivity, it has been unclear if there is any association between the local clustering of genesets and their prediction performance using SL. Here we show that the performance of SL, across networks and types of prediction tasks, is highly correlated with local network clustering of the genes of interest (Fig. 5 and Fig. S9). This result substantiates SL as an approach that can accurately predict gene attributes by taking advantage of local network connectivity.

While being accurate, training a supervised-learning model on the adjacency matrix (SL-A) can take some computational time and resources as the size of the molecular network increases, thus considerably differing in speed for, say, STRING-EXP (14,089 nodes and 141,629 unweighted edges) and GIANT-TN (25,689 nodes and 38,904,929 weighted edges). Worthy of note in this context is the recent excitement in deriving node embeddings for each node in a network, concisely encoding its connectivity to all other nodes, and using them as features in SL algorithms for node classification [60,69–74]. Although we show that SL-A markedly outperforms supervised-learning on the embedding matrix (SL-E; Fig. 6), the unique characteristics of SL-E methods call for further exploration. For instance, the greatly reduced number of features allows SL-E methods to be more readily applicable to classifiers more complex than logistic regression, such as deep neural networks (DNNs), which are typically ill-suited for problems where the number of features is much greater than the number of training examples. Further, since the reduced number of features allow SL-E methods to be trained orders of magnitude faster than SL-A or supervised-learning on the influence matrix (SL-I), they can be easily incorporated into ensemble learning models, which combine the results from many shallow learning algorithms. Akin to LP [38,75,76], node embeddings also offer a convenient route to incorporating multiple networks into SL approaches. While methods such as SL-I and SL-A may require concatenating the original networks or integrating them into a single network before learning, recent work has shown that SL-E-based methods can embed information from multiple molecular/heterogeneous networks and learn gene classifiers in tandem [77–85]. However, none of these studies have compared the variety of SL-E methods to learning directly on the adjacency matrix. Given our finding here that SL-A greatly outperforms SL-E for function, disease and trait prediction, we advice and urge that every new SL-E methods should be compared to SL-A for network-based gene classification.

In past work, SL methods for gene classification have mostly relied on hand-crafting features from graph-theory metrics, such as degree and centrality measures, or combining metrics to expand the feature set, resulting in a feature set size of ~30 or less [86,87]. We do not include a comparison to these types of methods in this study because predicting genes to functions or diseases based on generic network metrics such as high degree does not capture anything unique about specific functions or diseases. On the other hand, SL models with individual genes as features contain information biologically relevant to the specific prediction task [88,89].

Critical to all these conclusions is the rigorous preparation of diverse, specific prediction tasks and the choice of meaningful validation schemes and evaluation metrics. Temporal holdout and study-bias holdout validations help faithfully capture the performance of the computational methods when a researcher uses them to prioritize novel uncharacterized genes in existing molecular networks for experimental validation based on a handful of currently known genes. Although we provide all the results for the auROC metric in Supplemental Materials for completion (Figs. S4, S5, and S8), we base our conclusions on metrics driven by precision: auPRC and P@topK. While auROC is still commonly used in genomics, it is ill-suited to most biological prediction tasks including gene classification since they are highly imbalanced problems, with negative examples far outnumbering positive examples [90]. Optimizing for precision-based metrics, on the other hand, helps control for false-positives among the top candidates [91], thus making them pertinent to computational gene classification as a viable way of providing a list of candidate genes for further study. Accompanying the results in this manuscript, we are providing our comprehensive evaluation framework in the form of data – networks, prediction tasks, and evaluation splits – on Zenodo and the underlying code on Github to enable other researchers to not only reproduce our results but also to add new network-based gene classification methods for comparison. Together, the data and the code provides the community a systematic framework to conduct gene classification benchmarking studies. See “Availability of data and materials” for more information.

In conclusion, we have established that supervised-learning outperforms label-propagation for network-based gene classification across networks and prediction tasks (functions, diseases, and traits). We show that supervised-learning, in which every gene is its own feature, is able to capture network information just as well as label-propagation. Finally, we show that supervised-learning on the adjacency matrix demonstrably outperforms supervised-learning using node embeddings, and thus we strongly recommend that future work on using node embeddings for gene classification draws a comparison to using supervised-learning on the adjacency matrix.

## Additional files

**Additional file 1:** We provide supplemental material that contains detailed descriptions of the: molecular networks (Section 1.1), model selection and hyperparameter tuning (Section 1.2), processing steps and properties of the geneset-collections (Section 1.3), validation schemes (Section 1.4), evaluation metrics (Section 1.5), and results using P@topK, auROC, and 5FCV (Section 2).

## Acknowledgements

We are grateful to Daniel Marbach for making the GWAS data available. We thank members of the Krishnan Lab for valuable discussions and feedback on the manuscript.

## Author Contributions

AK, RL, and CAM conceived and designed the experiments; RL and CAM performed the experiments; RL, CAM, AY, KAJ, and AK processed the networks and geneset collections, and analyzed the data. RL, CAM, and AK wrote the manuscript. All authors read, edited, and approved the final manuscript.

## Funding

This work was primarily supported by US National Institutes of Health (NIH) grants R35 GM128765 to AK, and supported in part by MSU start-up funds to AK and MSU Engineering Distinguished Fellowship to AY.

## Availability of data and materials

The data used in this study, including the geneset-collections and networks, are freely available on Zenodo at https://zenodo.org/record/3352348. We note that KEGG and InBioMap data are available only from the original sources due to restrictive licences. A GitHub repository, GenePlexus, that contains code to reproduce the results in this study as well as add new gene classification methodsy is available at https://github.com/krishnanlab/GenePlexus.

## Notes

https://zenodo.org/record/3352348

https://github.com/krishnanlab/GenePlexus

