## Supplemental Material for "Supervised-learning is an accurate method for network-based gene classification"

### Section 1: Methods and Data

#### Section 1.1: Networks

The networks used in this study are BioGRID, STRING-EXP, InBioMap, GIANT-TN, and STRING. Detailed information about the network properties and sources can be seen in Table S1, with the network construction method and interaction type information coming from [1]. BioGRID (version 3.4.136) is a low-throughput network that includes both genetic interactions, as well as physical protein-protein interactions [2]. InBioMap (version 2016_09_12) is a high-throughput, scored network that contains physical protein-protein interactions as well as pathway database annotations incorporated as edges [3]. We used the “final-scores” as the edge weights. STRING (version 10.0) is a high-throughput, scored network that aggregates information from many data sources [4]. We used two different STRING networks. First, we used the “combined” network that directly includes database annotations, text-mining, ortholog information, co-expression, and physical protein interactions (referred to as “STRING” in this study). We also used a subset of edges in STRING that had just the “experiments” data, thus restricting the network to one constructed just from physical protein interactions in humans (referred to as “STRING-EXP” in this study). For both networks, we used the corresponding relationship scores as edge weights, after normalizing them to lie between 0 and 1. The GIANT-TN (version 1.0) network is the tissue-naïve network from GIANT [5], referred to as the “Global” network on the website, and is constructed from both low- and high-throughput data, and includes information from co-expression, non-protein sources, regulatory data, and physical protein-protein interactions. The GIANT-TN network is a fully connected, scored network. To add sparsity to the GIANT-TN network, we removed all edges with scores below 0.01 (equal to the prior the Bayesian model used to construct the network). It is worth noting here that the purpose of this study is not to compare networks against each other, but rather to determine the performance of SL methods vs LP methods on various types of networks.

**Table S1. Information on the molecular networks**. LT : low-throughput, HT : high-throughput, G : genetic, P : physical, DA : database annotations, CE : co-expression, NP : non-protein, R : regulation, CC : co-citation, O : orthologous.

| Network | Number of Genes | Number of Edges | Edge Density | Network Construction Method | Weighted | Interaction Type |
|---|---|---|---|---|---|---|
| BioGRID | 20,558 | 238,474 | 1.13e-3 | LT | No | G, P |
| STRING-EXP | 14,089 | 141,629 | 7.08e-4 | HT | Yes | P |
| InBioMap | 17,399 | 644,862 | 1.58e-3 | HT | Yes | P, DA |
| GIANT-TN | 25,689 | 38,904,929 | 1.92e-3 | LT, HT | Yes | CE, NP, P, R |
| STRING | 17,352 | 3,640,737 | 7.20e-3 | HT | Yes | CC, CE, O, DA, P |

### Section 1.2: Model Selection and Hyperparameter Tuning

The restart hyperparameter $\alpha$ used in generating an influence matrix was determined by doing a grid search over values between 0.55 and 0.95 in 0.1 steps for all networks and geneset collections, optimizing for auPRC using label propagation (Fig. S1). In general, there was not a strong dependence on $\alpha$. It can be seen in Figure S1 that a higher restart probability resulted in marginally better performance for the larger networks (STRING and GLOBAL), whereas as a smaller restart probability led to nominally better performance for the smaller networks such as BioGRID. In this study, we used $\alpha = 0.85$ for every geneset-collection–network combination, as $\alpha = 0.85$ offered good performance and had low variance. This $\alpha$ was used for both LP-I and SL-I. We stress that the tuning of $\alpha$ was never done for SL-I, and thus, our finding that SL methods generally outperform LP methods is not biased by this parameter tuning.

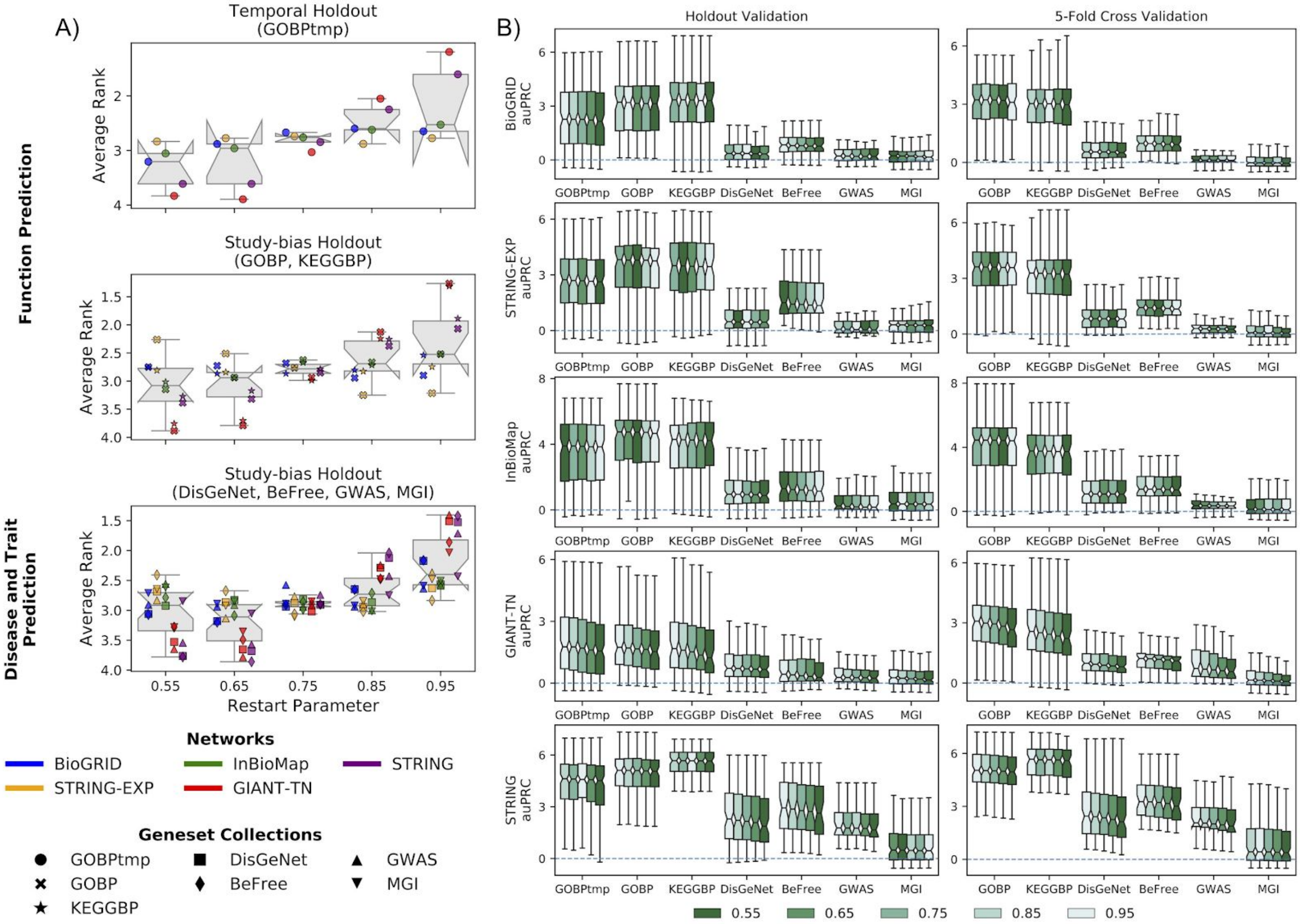


**Fig. S1. Tuning the restart probability hyperparameter for label propagation.** A) Each point in each boxplot represents the average rank for a geneset-collection–network combination, where the five restart probabilities that were tried were ranked in terms of performance (auPRC) for each geneset in a geneset-collection using the standard competition ranking. A restart probability of 0.85 was chosen for this study as it resulted in good overall performance as well as low variance in performance for the different genset-collection–network combinations. B) The performance for each individual geneset-collection–network combination is compared across the five restart probabilities: 0.55, 0.65, 0.75, 0.85 and 0.95. The methods are ranked by median value of auRPC with the highest scoring method on the left. There is no strong dependence of auRPC on the restart probability.

Model selection of the supervised learning classifier was done by comparing four popular classifiers that are implemented in Python package *Scikit Learn* [6]. To determine the best supervised learning classifier, we compared their performance over every geneset-collection–network combination using their default hyperparameters in version 0.19 of Scikit Learn (Fig. S2). Logistic regression with L2 regularization is marginally better than linear support vector machines and both these classifiers outperform random forest and logistic regression with L1 regularization.

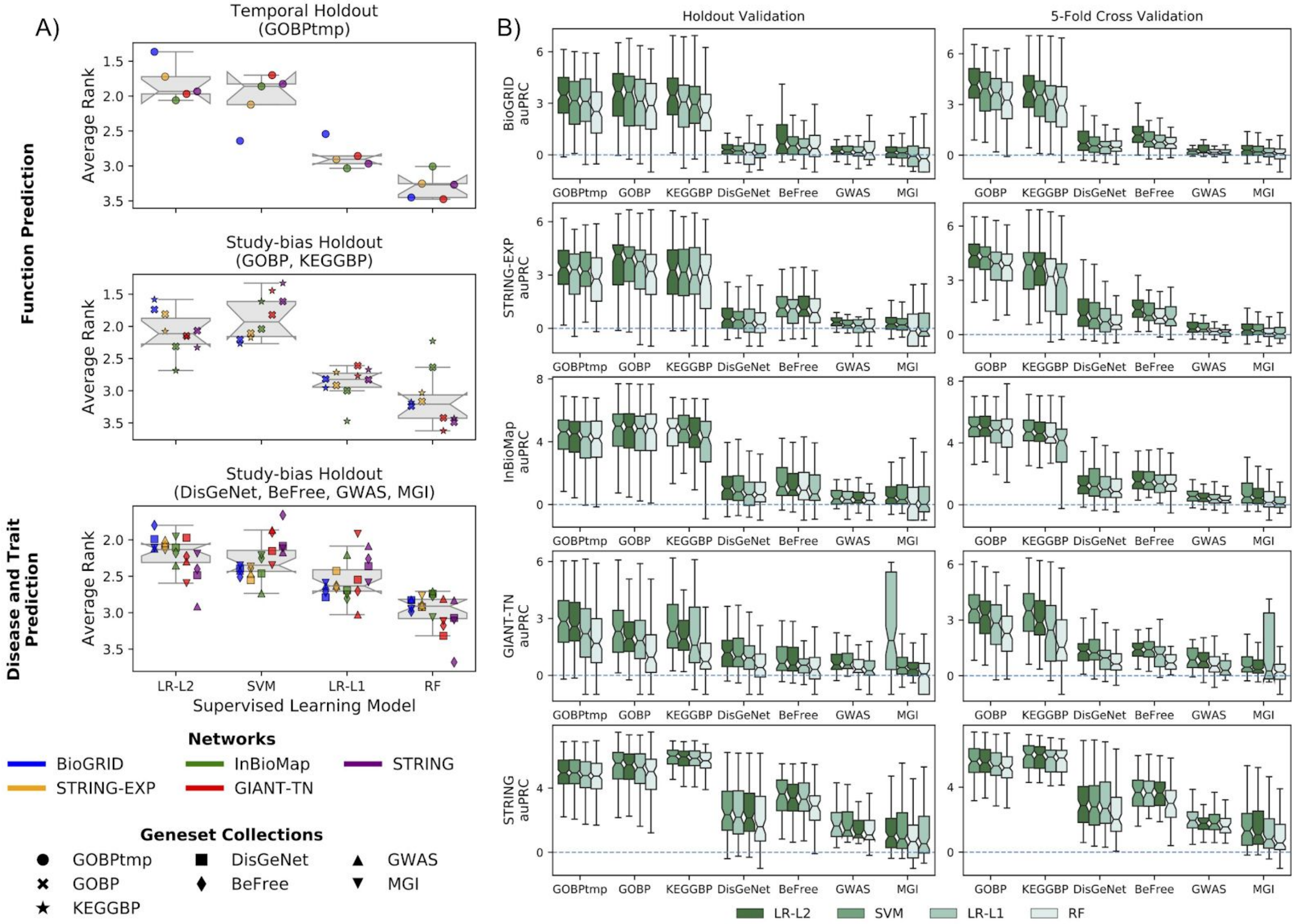


**Fig. S2. Comparison of classifiers for supervised learning.** A) Each point in each boxplot represents the average rank for a geneset-collection–network combination, where the four classifiers were ranked by the auPRC for each geneset in a geneset-collection using the standard competition ranking. Logistic regression with L2 regularization (LR-L2) was chosen as the classifier for supervised learning as it had slightly better overall performance than a linear support vector machine (SVM). B) The auRPC for each individual geneset-collection–network combination is compared across the four supervised learning classifiers: logistic-regression with L1 regularization (LR-L1), LR-L2, SVM, and a random forest (RF). The classifiers are ranked by median value with the best performing one on the left.

For the model selection of the embedding technique, we chose *node2vec* [7] because its competitive performance and ease of use [8]. The following hyperparameters were tuned based on the aggregated performance across all geneset-collections–network combinations: $p$ - the breadth first search parameter, $q$ - the depth first search parameter, $d$ - embedding size, $r$ - number of walks per node, $l$ - walk length, and $k$ - context window size. Since $p$ and $q$ are coupled, we performed a grid search for these two parameters leaving all others constant. Each of the other hyperparameters was tuned by leaving the rest at their default values as described in the original *node2vec* publication. This procedure resulted in the following hyperparameter values – $p = 0.1$, $q = 0.1$, $d = 512$, $k = 8$, $l = 120$, and $r = 10$ – which were used for every geneset-collection–network combination.

### Section 1.3: Geneset-collections

The geneset-collections used in this work are from the Gene Ontology (from version 2 of MyGene.info API with data retrieved on 2018-05-18, GOBPtmp, GOBP) [9–12], Kyoto Encyclopedia of Genes and Genomes (from version 6.1 of MSigDB, KEGGBP) [13–15], DisGeNet (version 5.0, DisGeNet, BeFree) [16,17], GWAS from a community challenge at https://www.synapse.org/#!Synapse:syn11944948 [18], and the Mouse Gene Informatics database (data retrieved on 2018-10-01, MGI) [19].

**Pre-processing genesets based on specificity, redundancy, and multi-functionality**

Each of these six geneset-collections contained anywhere from about a hundred to tens of thousands of genesets (Table S2) that varied widely in specificity and redundancy. The first pre-processing step we did after downloading the data was to convert the original gene/protein IDs to entrez gene IDs, which was done using gene ID conversions found in MyGene.info [11,12]. If the original ID mapped to more than one entrez ID, all of them were included for further analysis. Next, whenever applicable, annotations to genesets corresponding to terms in a curated ontology were propagated along the *is_a* and *part_of* relationships to ancestor terms in the corresponding ontologies: Gene Ontology [10] for GOBP, Disease Ontology [20] for DisGeNet and BeFree, and Mammalian Phenotype Ontology [21] for MGI.

Subsequent preprocessing steps were designed to ensure that the final set of genesets from each source are specific, largely non-overlapping, and not driven by multi-attribute genes.

***Specificity***: To select specific biologically-meaningful genesets in each collection, we sorted all the genesets in a collection from the largest to smallest based on the number of annotated genes (geneset size), manually examined their descriptions, and chose a size threshold that roughly separated large, generic genesets from the smaller, specific ones. This threshold was 200 for GOBP and KEGGBP, 300 for MGI, 400 for BeFree, 500 for GWAS and GOBPtmp, and 600 for DisGeNet.

***Redundancy***: To remove redundant genesets within a collection, first, we calculated the Jaccard index ($|A \cap B| / |A \cup B|$) and the overlap index ($|A \cap B| / min(|A|, |B|)$) between all pairs of genesets (with $A$ and $B$ representing the sets of genes annotated to the genesets). Then, we built a graph with the genesets as the nodes, and added edges between genesets pairs if their Jaccard index was >0.5 *and* their overlap index was >0.7. The geneset graph constructed in this manner contained many connected components, each representing a set of highly overlapping genesets. Finally, we used the following procedure to pick representative genesets within each component: a) calculate a score for each geneset equal to the sum of the proportions of genes in other linked genesets that are contained within it (higher this score, more representative that geneset is), b) create a sorted list of all the genesets in decreasing order of this score, and c) pick the first geneset in the list, remove every subsequent geneset that is connected to it in the graph, and repeat this step until the sorted list is empty. This procedure resulted in a reasonable number of non-redundant genesets within each collection. The same Jaccard and overlap thresholds were used for all collections except MGI. For MGI, since an overlap cutoff of 0.7 still resulted in thousands of genesets, it was further lowered to 0.5.

***Multi-attribute genes***: Given the set of largely non-overlapping genesets in a collection, individual genes were removed from all genesets if they appeared in more than 10 genesets in that collection. This step ensures that the evaluations are not biased by multi-attribute genes that can potentially be easily predicted in a non-specific manner [22].

We also note that we did not include the cellular component (CC) or molecular function (MF) classes of the gene ontology as part of the function classification tasks because two genes that are annotated to the same CC or MF need not be related to each other functionally.

**Table S2. Information on the geneset-collections**. The last four columns reflect the fact each geneset-collection is slightly different for every network and these values are presented as either a range, a median value, or number of genes in a union across all networks used in this study.

| Geneset Collection | Number of Genesets From Original Data | Number of Genesets After Redundant Genesets Removed | Number of Genesets After Holdout Preprocessing | Geneset Sizes | Median Geneset Size | Number of Genes from Union of all Genesets |
|---|---|---|---|---|---|---|
| GOBPtmp | 11,574 to 754* | 166 | (115, 160) | (27, 452) | 174 | 9464 |
| GOBP | 11,574 | 313 | (84, 96) | (20, 181) | 76 | 5301 |
| KEGGBP | 149 | 138 | (63, 74) | (24, 181) | 51 | 3454 |
| DisGeNet | 4030 | 334 | (89, 104) | (21, 368) | 67 | 4689 |
| BeFree | 2891 | 207 | (49, 57) | (20, 223) | 80 | 2692 |
| GWAS | 169 | 74 | (30, 37) | (24, 431) | 94 | 2134 |
| MGI | 10,264 | 492 | (90, 121) | (20, 132) | 41 | 2716 |

* The GOBP temporal holdout step had an extra initial preprocessing step to make sure there were at least ten genes in the training and testing sets.

#### Calculating the network properties of the genesets

To determine how the performance of a given geneset depends on the network, for each geneset we calculated three different properties:

1) For a given geneset, $T$, the number of genes annotated it is given by $|T|$.
2) For a given geneset, $T$, the edge density, $D_T$, is given by

$$D_T = \sum_{(u,v)\in T} W_{uv} / (|T| * (|T| - 1)/2)\,, \qquad \text{(eqn. S1)}$$

where $W_{uv}$ is the edge weight between genes $u$ and $v$. The edge density is a measure of how tightly connected the geneset is within itself.

3) For a given geneset, $T$, the segregation, $S_T$, is given by

$$S_T = \sum_{(u,v)\in T} W_{uv} / \sum_{u\in T,\, t\in V} W_{ut}\,. \qquad \text{(eqn. S2)}$$

Segregation is a measure of how isolated the geneset is from the rest of the network.

The three geneset properties are shown for all geneset-collection–network combinations in Fig. S3. In general, there is little difference in the number of genes across the different prediction tasks (i.e. function, disease and trait), except for GOBPtmp, which has the largest number of genes due to the fact the genesets need to be larger to have enough with at least 10 testing genes. Edge density and segregation are highest for the function genesets (GOBPtmp, GOBP, KEGGBP) and lowest for the disease and trait genesets (DisGeNet, BeFree, GWAS, MGI).

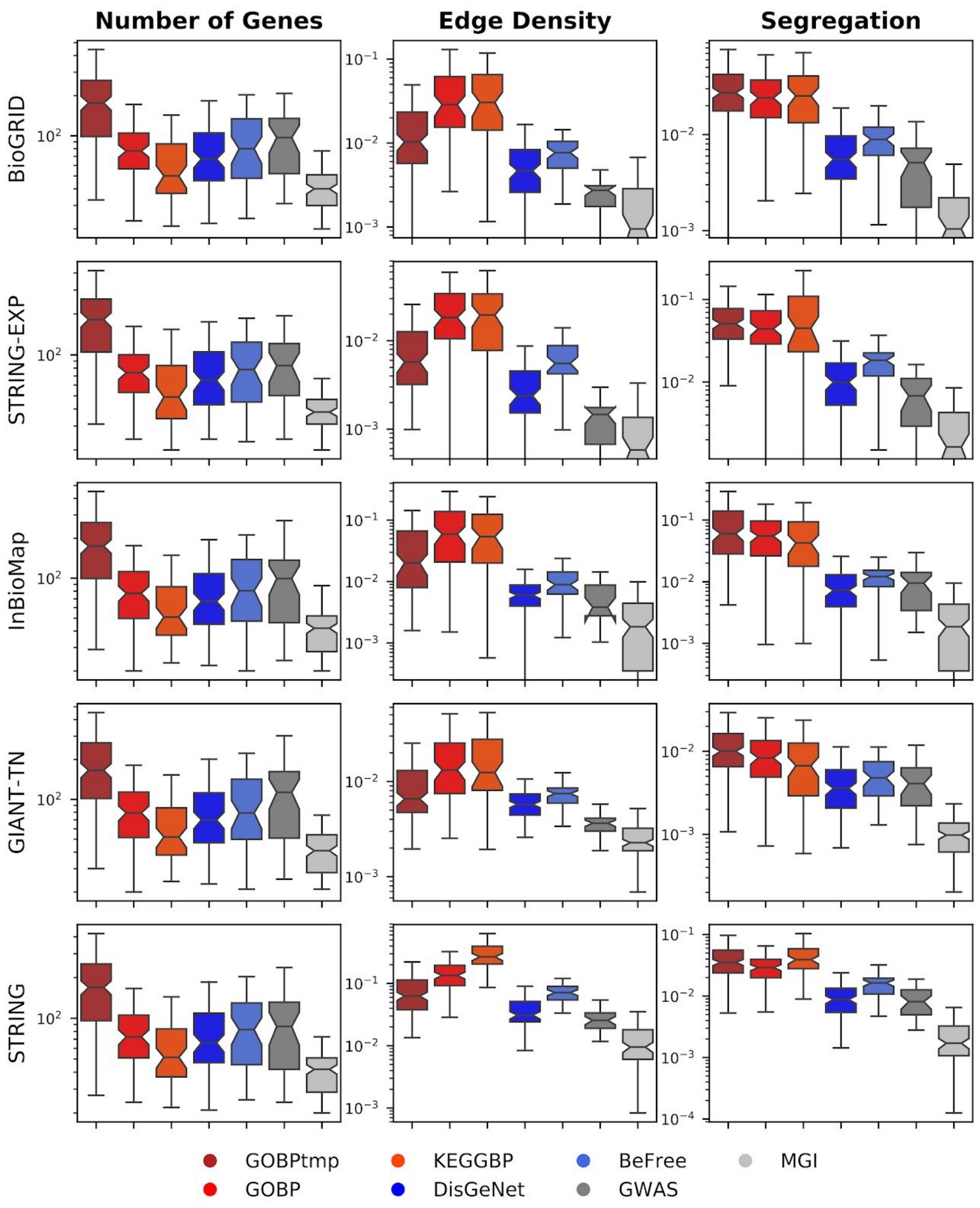


**Fig. S3. Network properties for the different geneset-collections.** The geneset-collections can be broken up into three prediction tasks; function (GOBPtmp, GOBP, KEGGBP; reds), disease (DiGeNet, BeFree; blues) and trait (GWAS, MGI; greys). In general, there is little difference in the number of genes across the different type prediction tasks (i.e. function, disease and trait), except for GOBPtmp which has the largest number of genes due to the fact the genesets need to be larger to have enough with at least 10 testing genes. Edge density and segregation are highest for the function genesets (GOBPtmp, GOBP, KEGGBP) and lowest for the disease and trait genesets (DisGeNet, BeFree, GWAS, MGI).

### Section 1.4: Validation schemes

We used three different validation schemes to evaluate gene classification.

#### Temporal holdout validation

Temporal holdout is the most stringent evaluation scheme for gene classification since it mimics the practical scenario of using current knowledge to predict the future. Since Gene Ontology was the only source with clear date-stamps for all its annotations, temporal holdout was applied only to the GOBP geneset-collection. Since the goal of this study is to use relatively recent and widely-used molecular networks, as this would reflect how these models would be deployed in practice, we chose a temporal cutoff point of Jan 1st, 2017. Then, for each geneset-collection, genes that only had an annotation to any geneset in the collection after 2017-01-01 were assigned to the testing set and the remaining genes were assigned to the training set. Since this resulted in the testing set having far fewer genes than the testing set for the other validation schemes, we made the following minor modifications to the geneset pre-processing procedure: GOBP geneset-collection was first filtered to remove any geneset with fewer than ten training genes or had fewer than ten testing genes based on the temporal split and the specificity threshold (maximum number of genes annotated to a geneset) was increased from 200 to 500. Redundancy filtering and multi-attribute gene filtering were unchanged. As noted in Section 1.1, from each network resource considered in this study, we chose the most recent version of the network that was released before 2017-01-01 to ensure no data leakage. Finally, genes were removed from genesets if they were not present in a given network, genesets with fewer than ten training genes or fewer than ten testing genes were filtered out, and the remaining genesets were used to perform the temporal holdout validation.

#### Study-bias holdout validation

The goal of study-bias validation is to evaluate the scenario that is close to the real-world situation of learning from well-characterized genes to predict novel un(der)-characterized genes. Here, we defined study-bias for each gene as the number of articles in PubMed (http://www.pubmed.gov/) in which that gene was referenced in, as determined in the *gene2pubmed* file (downloaded on 2018-10-30) from the NCBI Gene database [23]. Using this definition, for each geneset-collection–network combination, we created training-testing splits in the following manner: Genes were removed from genesets if they were not present in the given network. Then, among the remaining genes, a gene was assigned to the training set if it was in the top two-thirds of the list of genes sorted by their PubMed count. The remaining genes were assigned to the testing set. Finally, genesets with fewer than ten training genes or fewer than ten testing genes were filtered out and the remaining genesets were used to perform the study-bias holdout validation.

#### Five-fold cross-validation

To ensure comparability, we performed 5-fold cross-validation using the same genesets that were used in study-bias holdout, splitting each geneset randomly into five approximately equal folds (with similar proportions of positive and negative examples) and, in rotation, using one fold as the testing set and the remaining four as the training set.

### Section 1.5: Evaluation Metrics

In this study, we present results in terms of two popular metrics, auPRC and auROC, as well as the precision of the top *K* ranked predictions (P@TopK). Since, each geneset-collection–network combination has a different number of positive examples (and, hence, different positive:negative proportions), we normalized auPRC and P@topK by the prior. Specifically, auPRC is given by:

$$auPRC = \log_2\left( \frac{auPRC_S}{prior} \right) \quad \text{(eqn. S3)}$$

where $auPRC_S$ is the standard area under the precision-recall curve, and the *prior* is *P*/(*P*+*N*) with *P* being the number of positive ground truth labels, and *N* being the number of negative ground truth labels. The log2 in eqn. S3 allows for the following interpretation: the number of 2-fold increases of the measured $auPRC_S$ over what is expected given the ground truth labels (e.g., a value of 1 indicates a 2-fold increase, a value of 2 indicates a 4-fold increase). Similarly, P@topK is given by:

$$P@topK = \log_2\left(\frac{TP_K}{K \times prior}\right) \quad \text{(eqn. S4)}$$

where $K$ is the number of top-predictions to consider, $TP_K$ is the number true-positives of the top-K predictions, and the *prior* is the same as in eqn. S3. We set $K$ to be the number of ground truth positives in the testing set. P@topK can be thought of as what is the 2-fold increase in the percent of the top-K predictions that were predicted true over the expected value. Of note, it is possible that $TP_k = 0$ if no true positive is captured within the first *K* predictions. This causes P@topK to become $-\infty$. To address this issue, we set such values to be the minimum score obtained across all predictions for that given geneset-collection–network combination.

Precision-based metrics – auPRC and P@topK – are more suitable than the more popular area under the receiver-operating characteristic curve (auROC) for two reasons. First, gene classification is a highly imbalanced problem with many more negative samples than positive samples, and auROC is ill-suited for imbalanced problems [24]. Second, precision can control for Type-1 error (false positives) [25]. Since the foremost reason for gene classification is to provide a list of candidate genes for further experimental study, it is more important to make sure the top predictions are as correct as possible, as opposed to ensuring that, on average, positive examples are ranked higher than negative examples. However, for completeness, we have provided auROC results in this Supplemental Material (Section 2).

### Section 2: Supplementary Results

#### Section 2.1: Compiled results from all validation schemes in terms of all evaluation metrics

In this section, we present results for the ranking analysis used in Fig. 2, significance test analysis used in Fig. 3, as well as the boxplots representations seen in Fig. 4, based on all evaluation metrics (auPRC, P@TopK, and auROC) as well as all validation schemes (temporal holdout, study-bias holdout and 5FCV) (Figs. S4 - S8). Additionally, in this section, we present the results of how the performance of SL-A and LP-I scale with the number of genes, edge density, and segregation for all networks used in this study (Fig. S9).

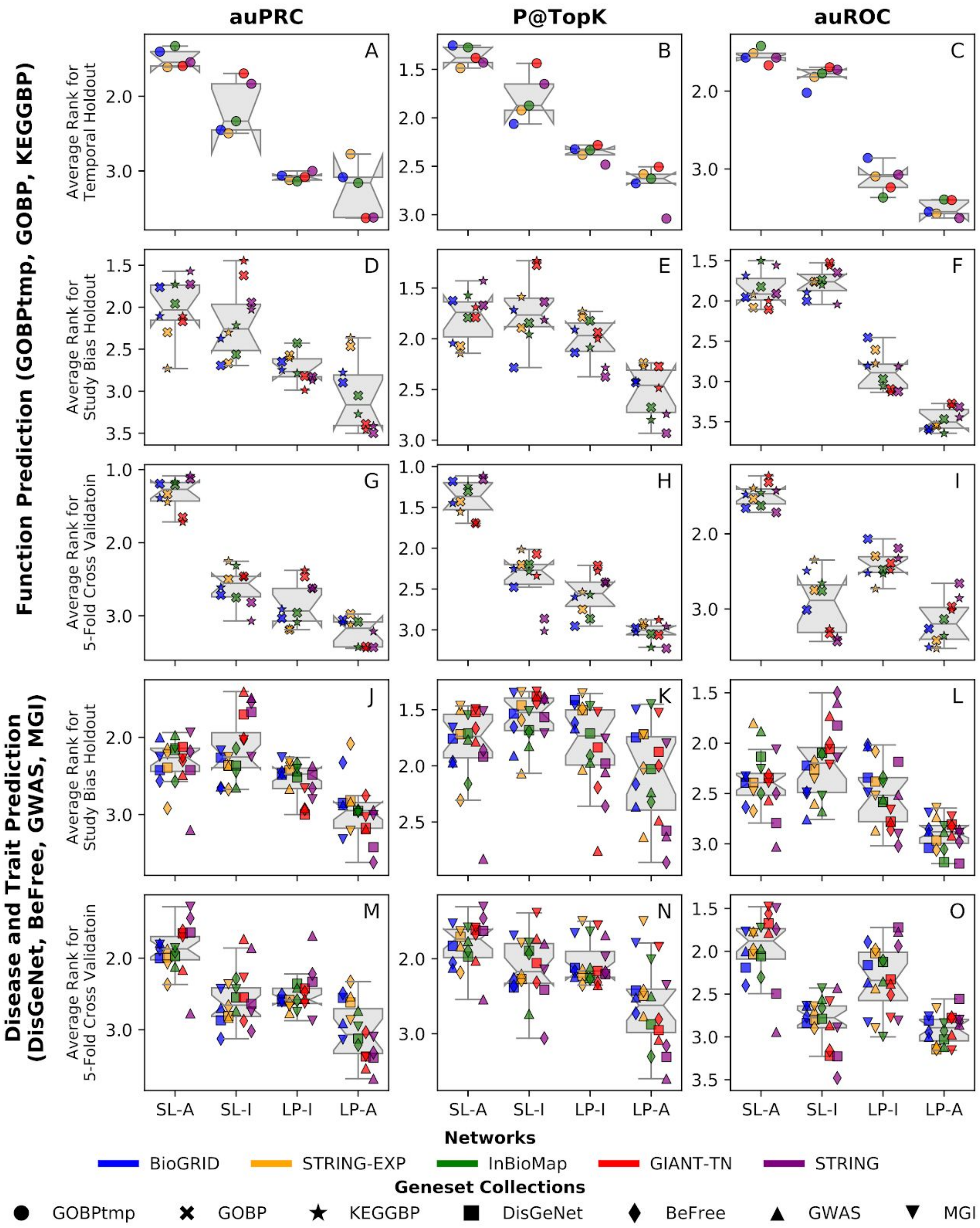


**Fig. S4. Average rank of the four methods for all evaluation metrics and validation schemes.** Each point in a boxplot represents the average rank for a geneset-collection–network combination, where the four methods were ranked in terms of performance for each geneset in a geneset-collection using the standard competition ranking. Different colors represent different networks and different marker shapes represent different geneset-collections.

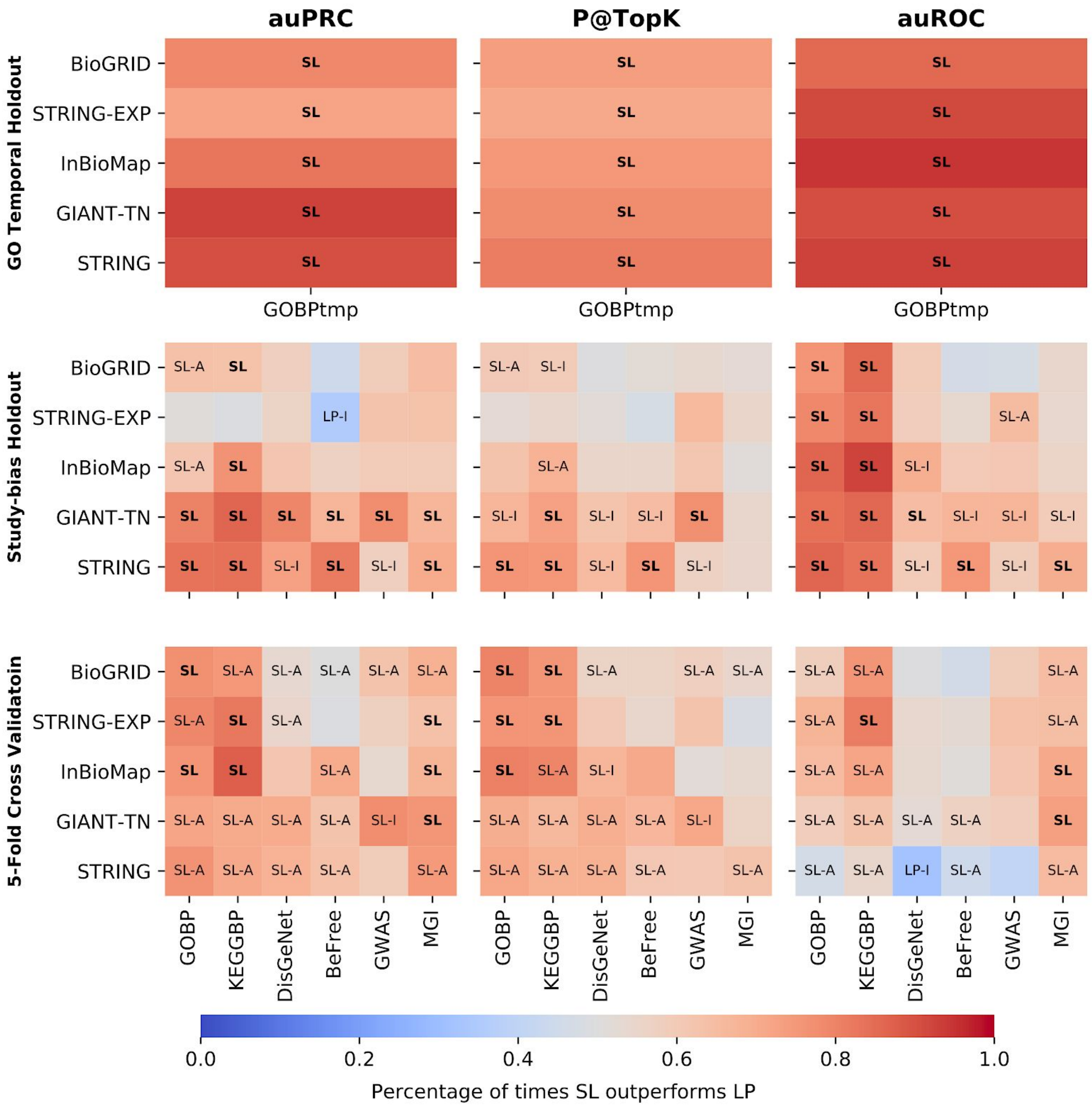


**Fig. S5. Testing for a statistically significant difference between SL and LP methods using all evaluation metrics and validation schemes.** For each network-geneset combination, each method is compared to the two methods from the other class (i.e. SL-A vs LP-I, SL-A vs LP-A, SL-I vs LP-I, SL-I vs LP-A). If a method was found to be significantly better than both methods from the other class (Wilcoxon ranked-sum test with an FDR threshold of 0.05), the cell is annotated with that method. If both models in that class were found to be significantly better than the two methods in the other class, the cell is annotated in bold with just the class. The color scale represents the fraction of genesets that were higher for the SL methods across all four comparisons. The first column uses GOBP temporal holdout, whereas the remaining 6 columns use study-bias holdout. B) SL methods show a statistically significant improvement over LP methods, especially for function prediction.

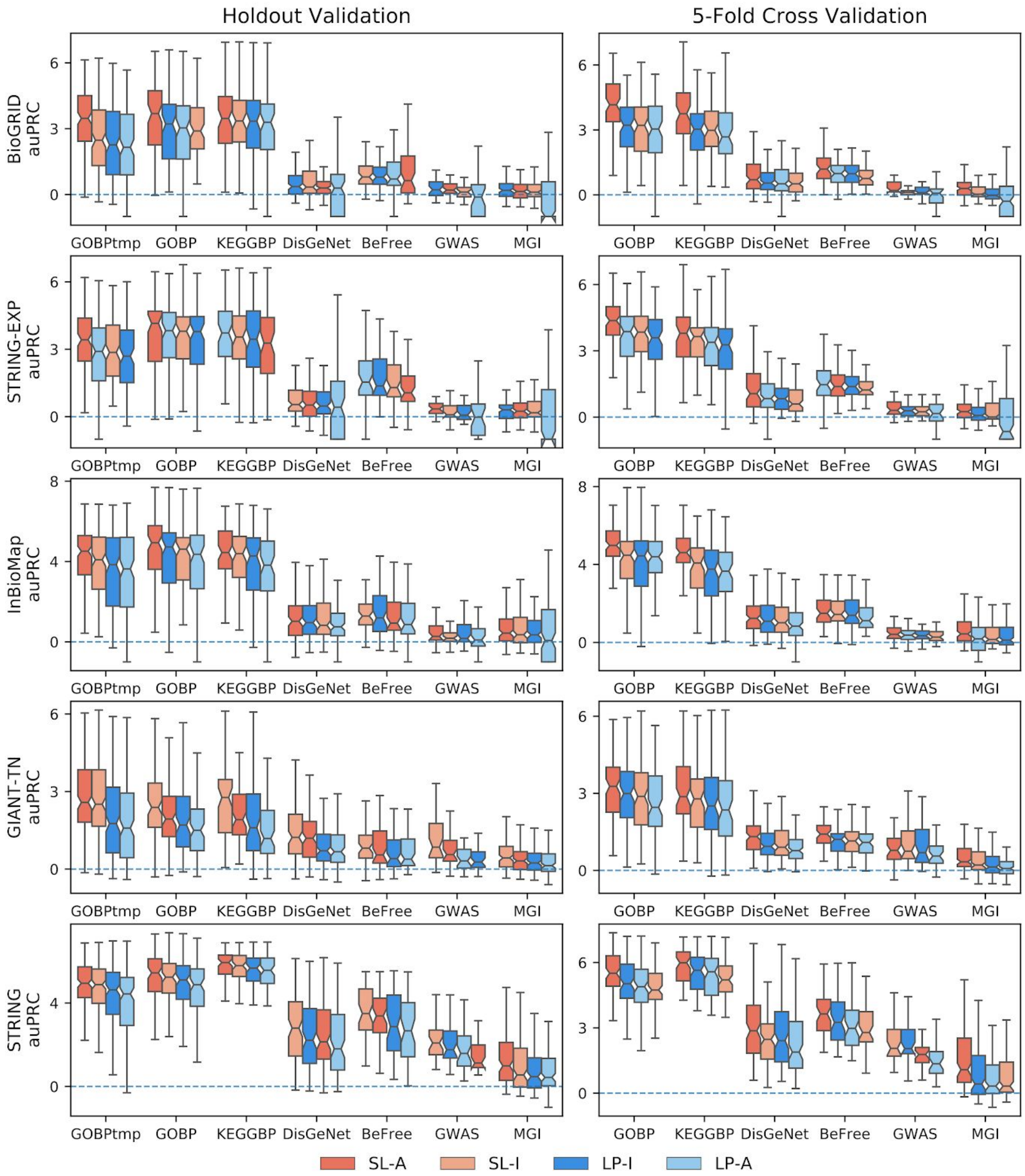


**Fig. S6. Boxplots for auPRC performance across all geneset-collection–network combinations.** The performance for each individual geneset-collection–network combination is compared across the four methods; SL-A (red), SL-I (light red), LP-I (blue), and LP-A (light blue). The methods are ranked by median value with the highest scoring method on the left. The first column contains temporal and study-bias holdout, and the second column is 5FCV. The scoring metric is auPRC.

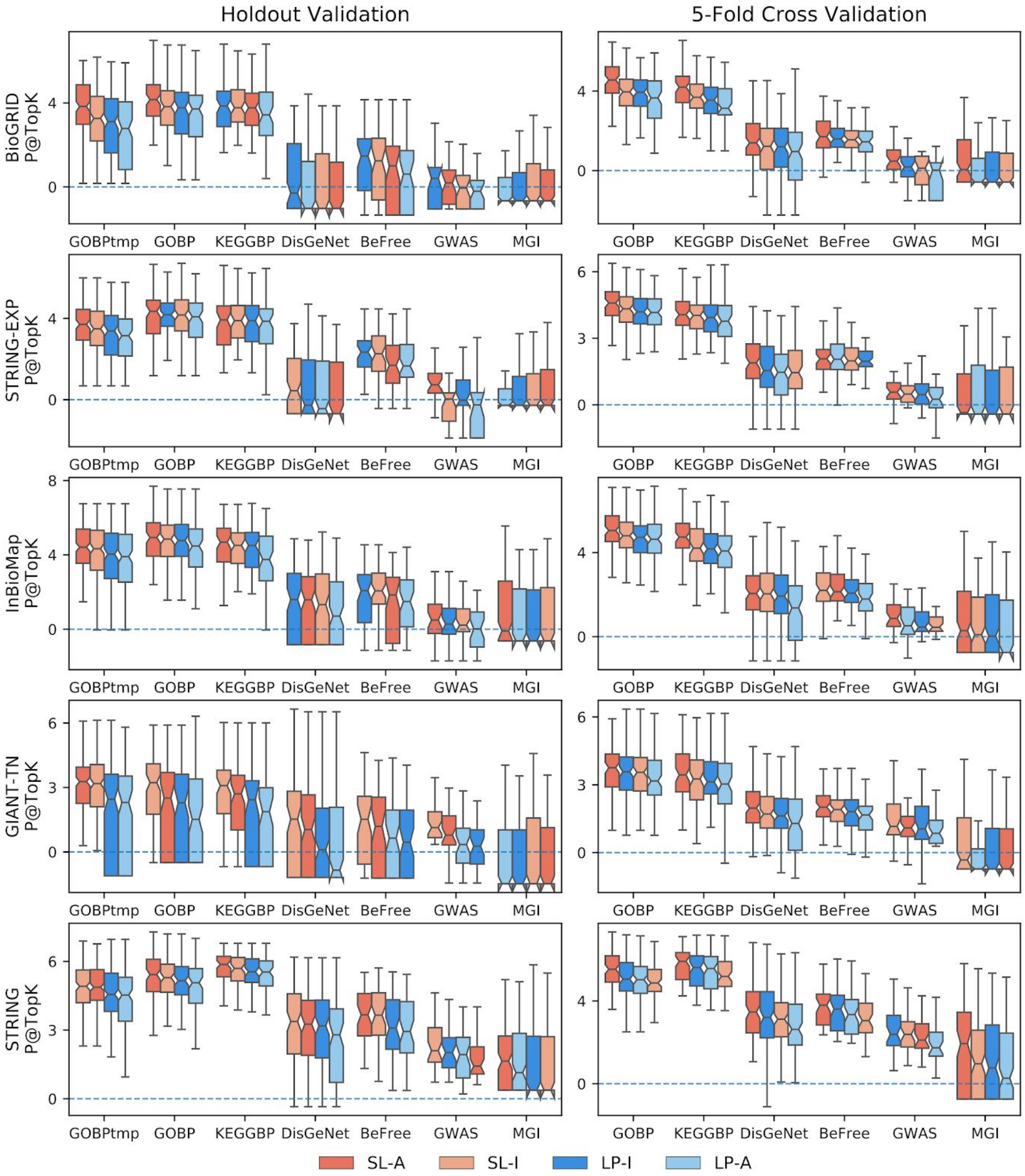


**Fig. S7. Boxplots for P@TopK performance across all geneset-collection–network combinations.** The performance for each individual geneset-collection–network combination is compared across the four methods; SL-A (red), SL-I (light red), LP-I (blue), and LP-A (light blue). The methods are ranked by median value with the highest scoring method on the left. The first column contains temporal and study-bias holdout, and the second column is 5FCV. The scoring metric is P@TopK.

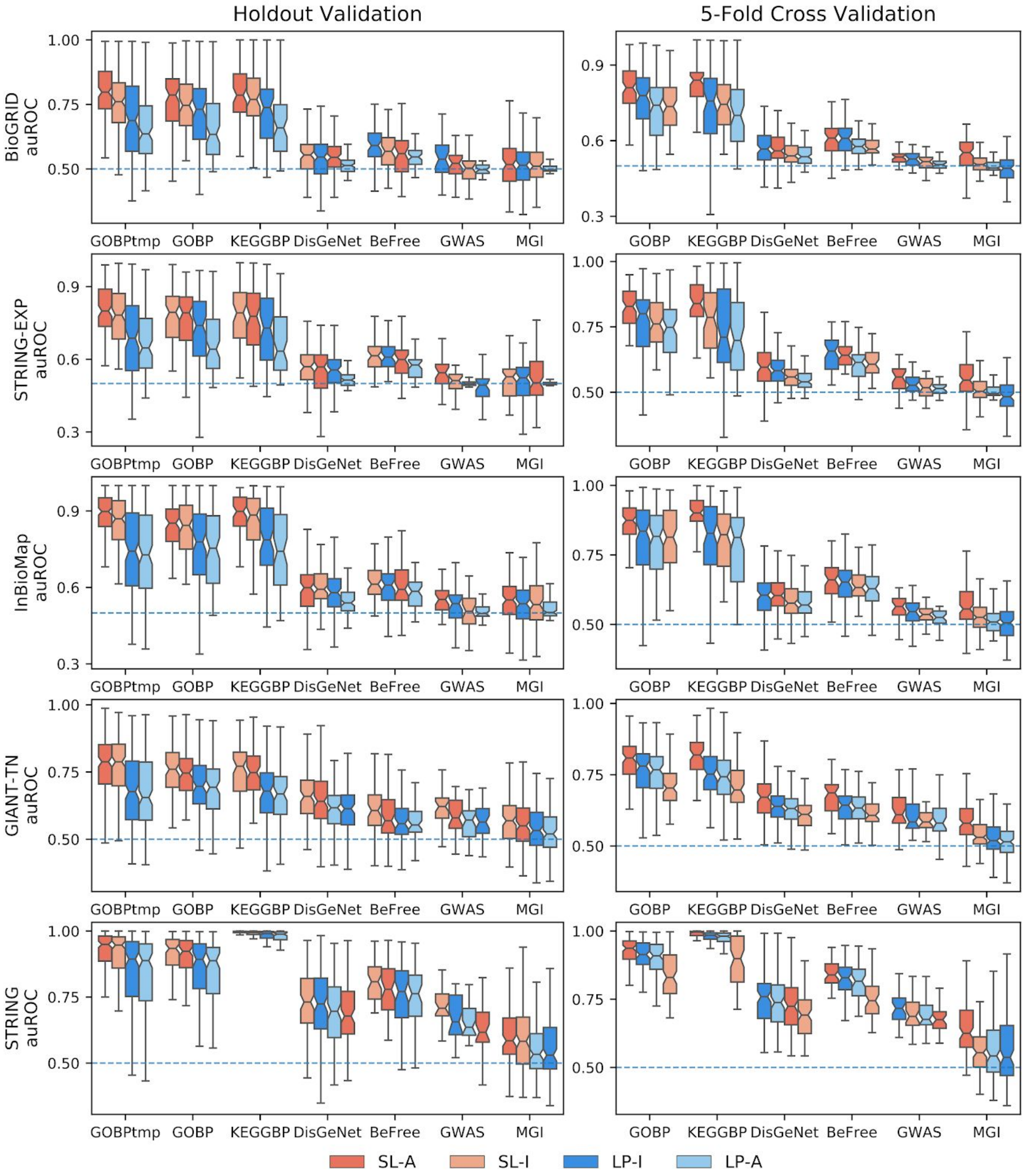


**Fig. S8. Boxplots for auROC performance across all geneset-collection–network combinations.** The performance for each individual geneset-collection–network combination is compared across the four methods; SL-A (red), SL-I (light red), LP-I (blue), and LP-A (light blue). The methods are ranked by median value with the highest scoring method on the left. The first column contains temporal and study-bias holdout, and the second column is 5FCV. The scoring metric is auROC.

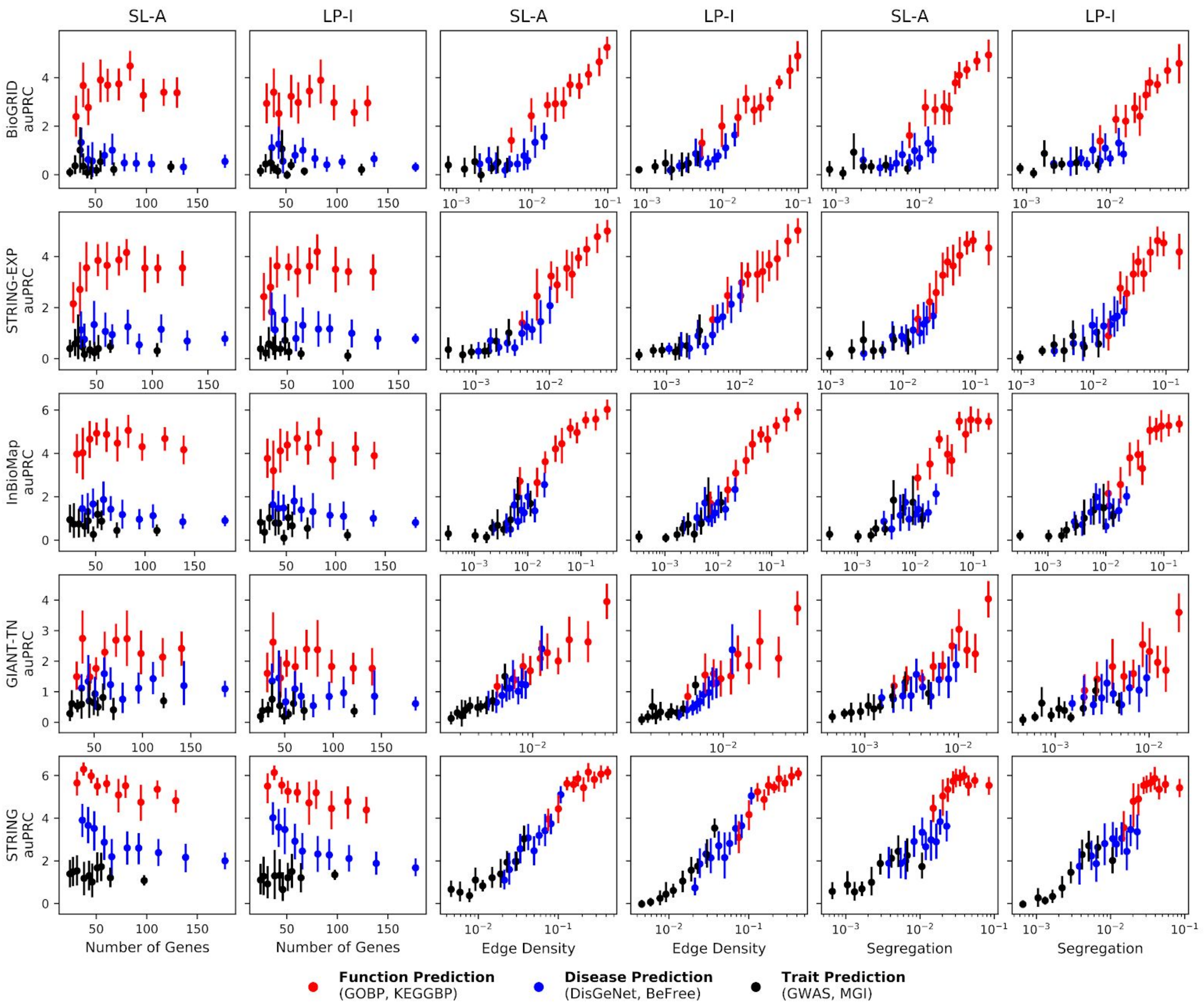


**Fig S9. Performance vs Network/Geneset properties for all networks.** SL-A is able to capture network information as efficiently as LP-I across all networks. There is no correlation between the number of genes in the geneset versus performance, but there is a strong correlation between the performance and the edge density as well as segregation. The different colored dots represent function genesets (red, GOBP and KEGGBP), disease genesets (blue, DIsGeNet and BeFree), and trait genesets (black, GWAS and MGI). The vertical line is the 95% confidence interval and the performance metric is auPRC.
